# Molecular basis of outer kinetochore assembly on CENP-T

**DOI:** 10.1101/071613

**Authors:** Pim J. Huis in ’t Veld, Sadasivam Jeganathan, Arsen Petrovic, Juliane John, Priyanka Singh, Florian Weissmann, Tanja Bange, Andrea Musacchio

## Abstract

Stable kinetochore-microtubule attachment is essential for cell division. It requires recruitment of outer kinetochore microtubule binders by centromere proteins C and T (CENP-C and CENP-T). To study the molecular requirements of kinetochore formation, we reconstituted the binding of the MIS12 and NDC80 outer kinetochore subcomplexes to CENP-C and CENP-T. Whereas CENP-C recruits a single MIS12:NDC80 complex, we show here that CENP-T binds one MIS12:NDC80 and two NDC80 complexes upon phosphorylation by the mitotic CDK1:Cyclin B complex at three distinct CENP-T sites. Visualization of reconstituted complexes by electron microscopy supports this model. Binding of CENP-C and CENP-T to MIS12 is competitive, and therefore CENP-C and CENP-T act in parallel to recruit two MIS12 and up to four NDC80 complexes. Our observations provide a molecular explanation for the stoichiometry of kinetochore components and its cell cycle regulation, and highlight how outer kinetochore modules bridge distances of well over 100 nm.

## Introduction

Accurate chromosome segregation in eukaryotes requires the coordinated action of hundreds of proteins. Subsets of these assemble on centromeric chromatin that is epigenetically specified by the enrichment of centromeric protein A (CENP-A), a variant of Histone H3 (Guse et al., 2011). These assemblies, named kinetochores, form the major point of attachment between centromeres and the mitotic or meiotic spindle and couple the force of depolymerizing microtubules to chromosome movement (Cheeseman, 2014). Kinetochores also function as signaling platforms for the spindle assembly checkpoint and delay the onset of chromosome segregation in the presence of erroneous chromosome-spindle attachments (Musacchio, 2015).

The affinity of the kinetochore for microtubules is predominantly mediated by the NDC80 complex (NDC80C), a heterotetramer with an approximately 50 nm long coiled coil region that separates the microtubule-binding calponin homology (CH) domains of the NDC80 and NUF2 subunits from the SPC24 and SPC25 subunits (Ciferri et al., 2005; Ciferri et al., 2008; Wei et al., 2007; Wei et al., 2005). NDC80C is essential to form stable regulated kinetochore-microtubule interactions and its localization at the outer kinetochore is thus a prerequisite for faithful chromosome segregation (Cheeseman et al., 2006; DeLuca et al., 2005; DeLuca et al., 2006). The recruitment of NDC80C requires the inner kinetochore proteins CENP-C and CENP-T (Carroll et al., 2010; Gascoigne et al., 2011; Hori et al., 2013; Kwon et al., 2007; Liu et al., 2006; Okada et al., 2006) as well as nuclear envelope breakdown and the activity of mitotic kinases (Gascoigne and Cheeseman, 2013).

Together with a set of other proteins, CENP-C and CENP-T are part of a larger protein complex at the inner kinetochore, also known as the constitutive centromere associated network (CCAN). Whereas CENP-C directly binds to CENP-A containing nucleosomes (Carroll et al., 2010) and has been proposes to act as a blueprint for further kinetochore assembly (Klare et al., 2015), CENP-T is integrated in a CENP-TWSX complex that requires its DNA-binding activity (Hori et al., 2008; Nishino et al., 2012), as well as interactions with the CCAN (Basilico et al., 2014; Carroll et al., 2010; Logsdon et al., 2015; Pekgoz Altunkaya et al., 2016; Samejima et al., 2015; Suzuki et al., 2015), to localize to the kinetochore.

Depletions of CENP-C or CENP-T, or prevention of cyclin-dependent kinase (CDK) phosphorylation of CENP-T all affect proper recruitment of MIS12C and NDC80C to the kinetochore (Gascoigne and Cheeseman, 2013; Gascoigne et al., 2011; Hori et al., 2013; Kim and Yu, 2015; Rago et al., 2015; Suzuki et al., 2015). Studying the respective contribution of CENP-C and CENP-T to outer kinetochore assembly, however, has been complicated by their interdependent localization and their distinct regulation by post-translational modifications. Experiments that targeted either CENP-C or CENP-T to an ectopic chromatin array uncoupled these requirements and demonstrated the ability of both pathways to recruit outer kinetochore components (Gascoigne et al., 2011). An important conclusion from these previous studies is that CENP-C and CENP-T recruit NDC80C in distinct ways (**Figure 1A**). The CENP-C-dependent axis, which is now understood in detail, relies on a direct interaction of CENP-C with the MIS12 complex (MIS12C) (Przewloka et al., 2011; Screpanti et al., 2011), which further recruits NDC80C via a direct interaction (Cheeseman et al., 2006; Petrovic et al., 2010).

**Figure 1.**
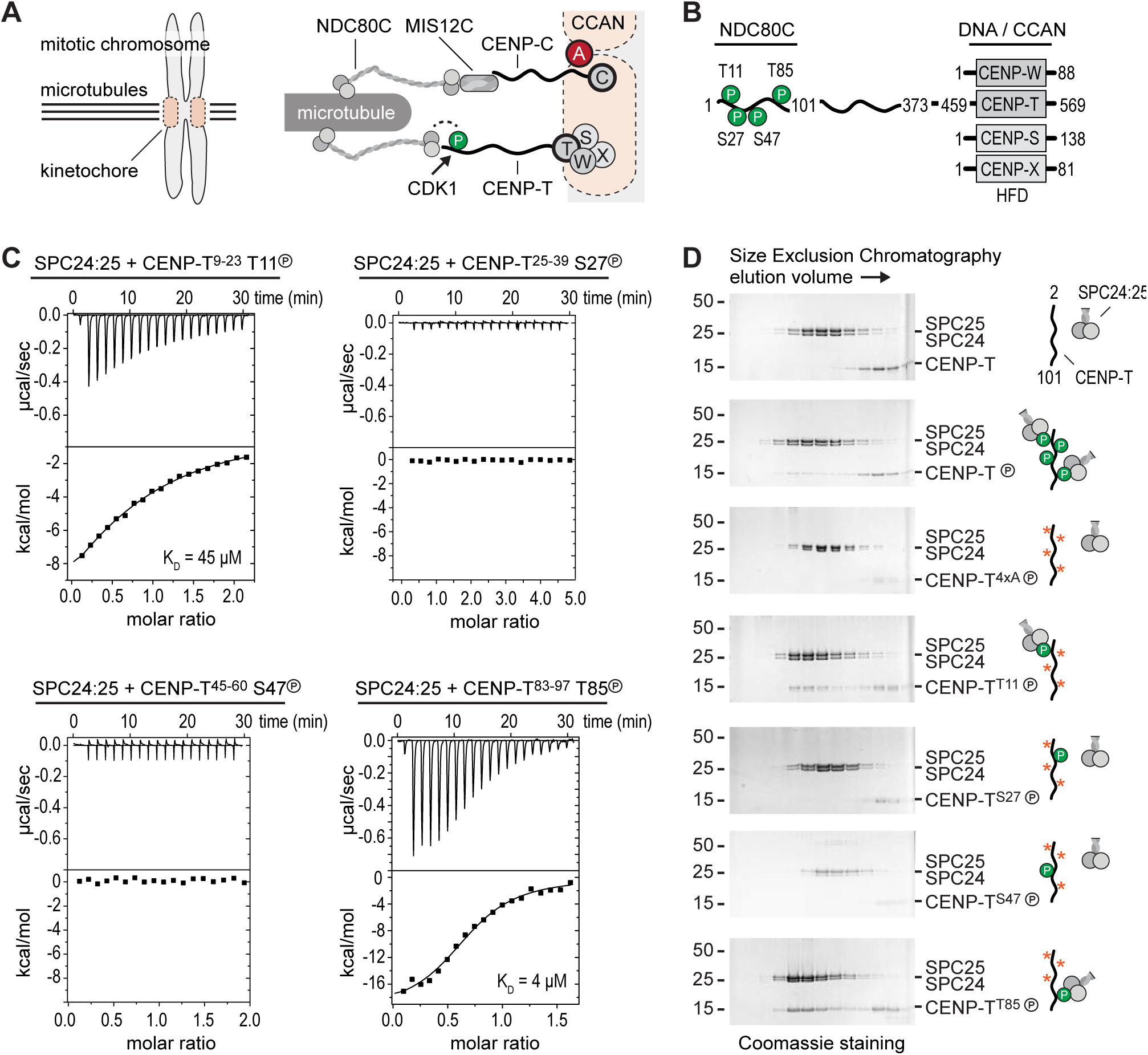
Phosphorylation of CENP-T^2-101^ at T11 or T85 is sufficient for the binding of SPC24:SPC25. (**A**) Schematic representation of CENP-C and CENP-T recruiting MIS12C:NDC80C and NDC80C. **(B**) Besides the histone-fold domain (HFD) at the carboxy terminus, CENP-T mainly consists of regions of compositional bias, rich in polar residues, and likely to be intrinsically disordered. Boundaries of CENP-T constructs used in this study and domains involved in the binding of NDC80C and DNA/CCAN are indicated. *In vitro* phosphorylated residues in CENP-T^1-101^ are marked by a P (see also Table 1). (**C**) The binding between SPC24:SPC25 and four CENP-T phosphopeptides was determined by isothermal titration calorimetry. The y-axis indicates kcal/mole of injectant. Dissociation constants between SPC24:SPC25 and phosphopeptides containing T11 and T85 were determined to be 45 µM and 4 µM respectively. (**D**) SDS-PAGE analysis of various CENP-T^2-101^ mutants that were incubated with SPC24:SPC25 and separated by analytical size-exclusion chromatography showing that phosphorylation of CENP-T at T11 or T85 is sufficient for the binding of SPC24:SPC25. Phosphorylation of T27 and S47 is dispensable. Red asterisks indicate mutated phosphorylation sites. See also figure 1 - figure supplement 1 for the entire dataset.

The CENP-T-dependent pathway, on the other hand, remains less well characterized. It is firmly established that CENP-T interacts directly with NDC80C, with (humans and chicken) or without (budding yeast) prior phosphorylation by CDK1:Cyclin B of sequence motifs in the CENP-T N-terminal region (Gascoigne et al., 2011; Malvezzi et al., 2013; Nishino et al., 2013; Schleiffer et al., 2012). Complicating the picture, however, CENP-T also contributes to kinetochore recruitment of MIS12C (Kim and Yu, 2015; Rago et al., 2015). This may appear surprising, because MIS12C and CENP-T bind to NDC80C in a competitive manner (Nishino et al., 2013; Schleiffer et al., 2012), suggesting that they can only recruit NDC80C independently from each other. Whether the interaction of CENP-T with MIS12C is direct, and compatible with CENP-T binding to NDC80C, is therefore currently unclear.

The ability of kinetochores to form multiple low-affinity linkages with microtubules is at the basis of popular models of kinetochore-microtubule attachment such as the Hill’s sleeve (Hill, 1985). It is also plausible that crucial regulatory aspects of kinetochore-microtubule attachment, such as the tension-dependent regulation required for correcting erroneous attachments or for stabilization of correct ones, depend on the effective number and distribution of linkages, as recently proposed in elegant modeling studies (Zaytsev et al., 2014). A detailed understanding of the stoichiometry of outer kinetochore composition and of its regulation is therefore crucial.

To address this question, we have embarked in an effort of biochemical reconstitution of kinetochores. This recently allowed us to report that the CENP-A nucleosome recruits two copies of the CCAN complex (Weir et al., 2016). In this study, we address instead the composition and regulation of the interactions between CCAN and the outer kinetochore, focusing on the interactions of CENP-T, MIS12C, and NDC80C. *In vitro* CDK1:Cyclin B phosphorylation reactions with systematically mutated CENP-T substrates revealed that the phosphorylation of CENP-T at Thr11 and Thr85 results in the recruitment of two NDC80Cs. We also demonstrate that CENP-T phosphorylated at Ser201 directly binds MIS12C:NDC80C. Our electron microscopy analysis demonstrates unequivocally that CENP-T binds one MIS12C and up to three NDC80Cs. Furthermore, we show that CENP-C and CENP-T are competitive MIS12C binders. Thus, we have reconstituted and visualized the molecular basis of CENP-T mediated assembly of the outer kinetochore and its phospho-regulation.

## Results

### Phosphorylation of CENP-T at T11 or T85 is sufficient to recruit SPC24:SPC25

To obtain insight into the molecular mechanism of CENP-T mediated recruitment of the human outer kinetochore, we set out to reconstitute this process using purified components. Previous studies identified multiple phosphorylation sites in CENP-T and showed that its phosphorylation contributes to the recruitment of NDC80C (Gascoigne and Cheeseman, 2013; Gascoigne et al., 2011; Kettenbach et al., 2011; Nishino et al., 2013; Rago et al., 2015). Confirming this, we identified numerous residues in CENP-T by mass spectrometry that were *in vitro* phosphorylated by CDK1:Cyclin B (**Table 1**). This included the CDK target sites Thr11, Thr27, Ser47, and Thr85 in the N-terminal region of CENP-T, which binds to the SPC24:SPC25 subunits of NDC80C (Malvezzi et al., 2013; Nishino et al., 2013) (**Figure 1B**). We obtained four short synthetic peptides that include phosphorylated forms of these residues and tested their binding with SPC24:SPC25 using isothermal titration calorimetry experiments. Peptides containing phosphorylated T11 or T85 showed a binding affinity of 45 µM and 4 µM respectively whereas peptides containing phosphorylated T27 or S47 did not show an interaction with SPC24:SPC25 (**Figure 1C**). Although the determined affinities are modest and unlikely to be representative for the binding affinities between full-length CENP-T and SPC24:SPC25, they distinguish *bona fide* binding sites from sites that do not interact.

**Table 1.**
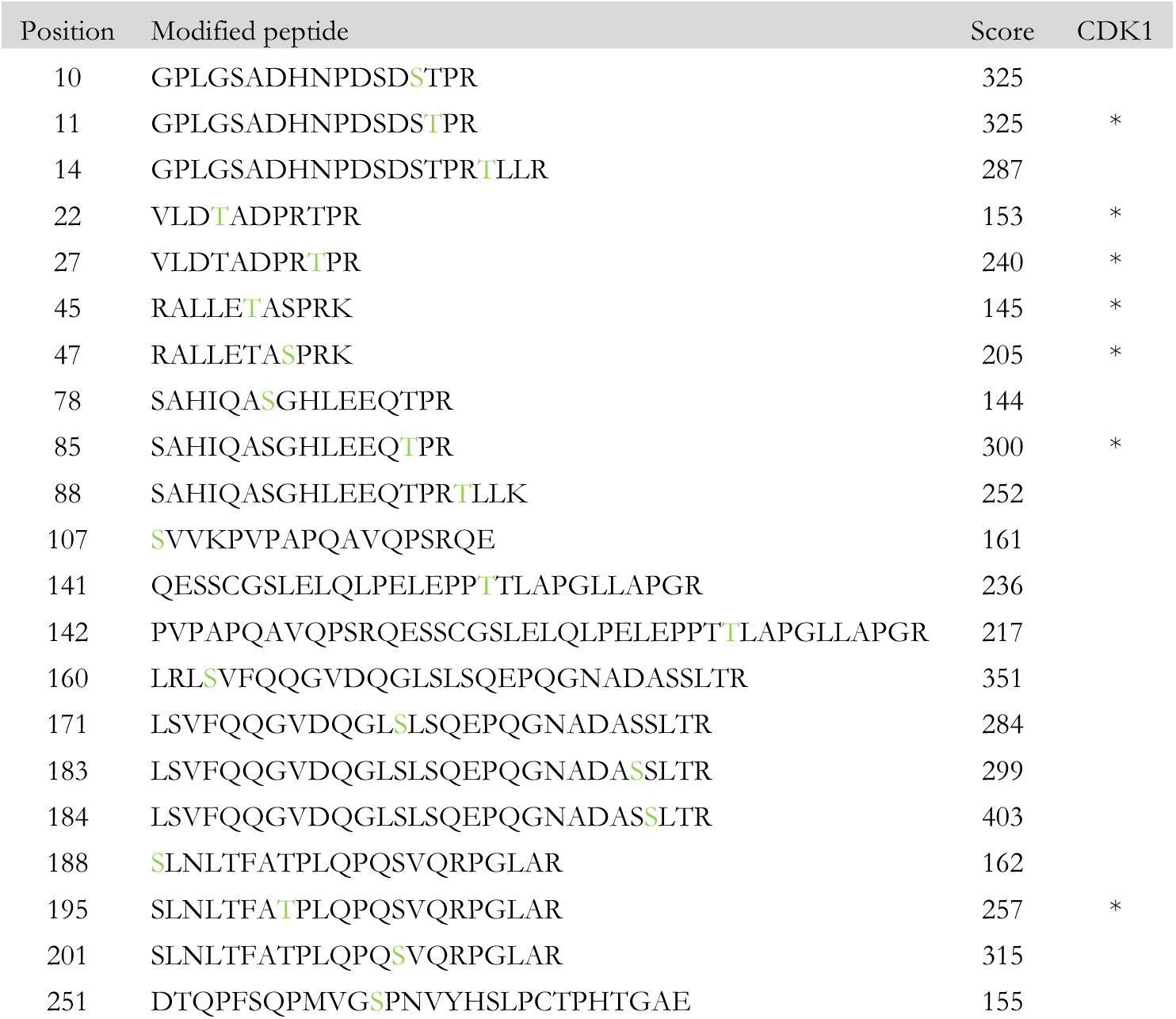
Phospho-sites on CENP-T^2-373^ *in vitro* phosphorylated by CDK1:Cyclin B. Identified phospho-peptides with a localization probability for the phospho-site > 90% and an Andromeda search engine score >140 are shown. All peptides have a posterior error probability (PEP) below 1x10^−30^. Modified sites are highlighted in green. Asterisks indicate CDK consensus sites (Xue et al., 2011).

Confirming these equilibrium binding experiments, the interaction between an N-terminal fragment of purified CENP-T and SPC24:SPC25 depended on the phosphorylation of CENP-T by CDK1:Cyclin B (**Figure 1D** and **Figure 1 - figure supplement 1**). We next generated a set of CENP-T^2-101^ constructs in which three out of the four CDK phosphorylation sites were mutated to Alanine and found that fragments retaining either Thr11 or Thr85 bound to SPC24:SPC25 in a phosphorylation dependent manner, whereas fragments retaining either Thr27 or Ser47, or having all four sites mutated to Alanine, did not bind SPC24:SPC25 (**Figure 1D** and **Figure 1 - figure supplement 1**). Phosphorylation of CENP-T at T11 or T85 is thus sufficient to bind SPC24:SPC25. These results are consistent with *in vivo* experiments showing that ectopic chromosome anchoring of a CENP-T^1-250^ fragment results in the recruitment of NDC80C when either Thr11 or Thr85 were mutated to Alanine, but not when both were mutated (Rago et al., 2015).

### CENP-T coordinates two NDC80 complexes in a single complex

The finding that phosphorylation of CENP-T at either Thr11 or Thr85 is sufficient to recruit SPC24:SPC25, both *in vivo* (Rago et al., 2015) and *in vitro* (**Figure 1**), raises the question whether phosphorylated CENP-T can bind two SPC24:SPC25 units simultaneously. This possibility is supported by the change in retention volume and the apparent stoichiometry of the CENP-T^2-101^:SPC24:SPC25 complex (**Figure 1D**). We set out to test the stoichiometry of CENP-T:SPC24:SPC25 complexes further using a CENP-T^2-373^ fragment that we C-terminally labeled with a fluorescent dye to specifically follow CENP-T during chromatography and SDS-PAGE (**Figure 2**). This is helpful because CENP-T^2-373^, which lacks Tryptophan and has only one Tyrosine, absorbs poorly at 280 nm. Fluorescently labeled CENP-T^2-373^ (hereafter CENP-T) was *in vitro* phosphorylated by CDK1:Cyclin B and subsequently incubated with a threefold molar excess of SPC24:SPC25. CENP-T was part of a broad peak with two SPC24:SPC25- containing species with approximate retention volumes of 1.2 and 1.35 ml (**Figure 2**, green traces). Consistent with the idea that the complex eluting at 1.2 mL contained two SPC24:SPC25 molecules per CENP-T, this species was not observed for CENP-T containing the T11A or the T85A mutation (**Figure 2**, orange traces). CENP-T that was mutated at both T11 and T85 or that was not phosphorylated, did not form a complex with SPC24:SPC25 (**Figure 2**, red and black traces).

**Figure 2.**
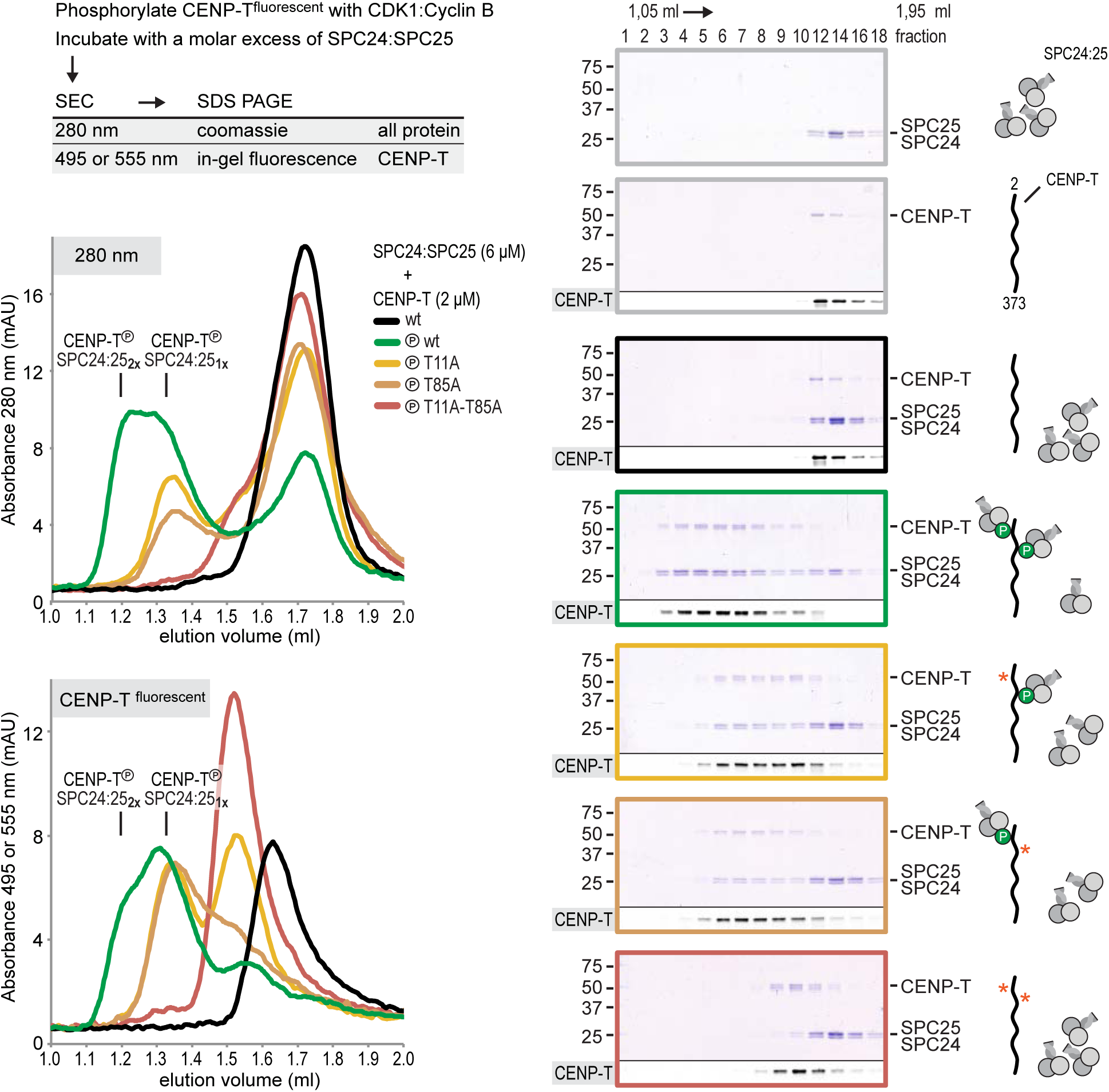
CENP-T phosphorylated at T11 and T85 binds two copies of SPC24:SPC25. Analytical size-exclusion chromatography (Superdex 200 5/150 increase) and SDS-PAGE show that CENP-T^2-373^ phosphorylated by CDK1:Cyclin B at positions T11 and T85 can bind two copies of SPC24:SPC25. CENP-T was fluorescently labeled with FAM (wt) or TMR (mutants) and monitored specifically during chromatography and after SDS-PAGE. Red asterisks indicate mutated phosphorylation sites.

Using a similar experimental setup, we determined that CENP-T also binds full-length NDC80Cs in a manner that depends on the phosphorylation of T11 and T85 (**Figure 3A** and **Figure 3 - figure supplement 1**). We next set out to visualize these assemblies directly by electron microscopy (EM) after low-angle metal shadowing. This technique enhances the contrast of the thin coiled-coil regions of NDC80C, enabling a characterization of the overall dimensions of complexes containing NDC80C. We inspected a number of NDC80Cs in detail (*n*=150). The coiled-coil of NDC80C spans 51 nm and the NDC80:NUF2 - SPC24:SPC25 end-to-end length is 62 nm (corrected for an estimated 3 nm shadow contribution at both ends) (**Figure 3B** and **Figure 3 - figure supplement 2**). This is slightly longer than the 57 nm of yeast NDC80C previously determined using the same technique (Wei et al., 2005). The difference might be explained by the use of full-length human NDC80C whereas the used yeast version lacked the N-terminal 100 residues of the NDC80p subunit, although interspecies or technical differences cannot be excluded. The low-resolution and the flexible appearance of the NDC80C prevent unambiguous assignment of the SPC24:SPC25 and the NDC80:NUF2 modules of the complexes, although a clear kink in the coiled coil, previously attributed to a sequence insertion in NDC80 (Ciferri et al., 2008) marks NDC80:NUF2 in a number of cases (**Figure 3B** and **Figure 3 - figure supplement 2**).

**Figure 3.**
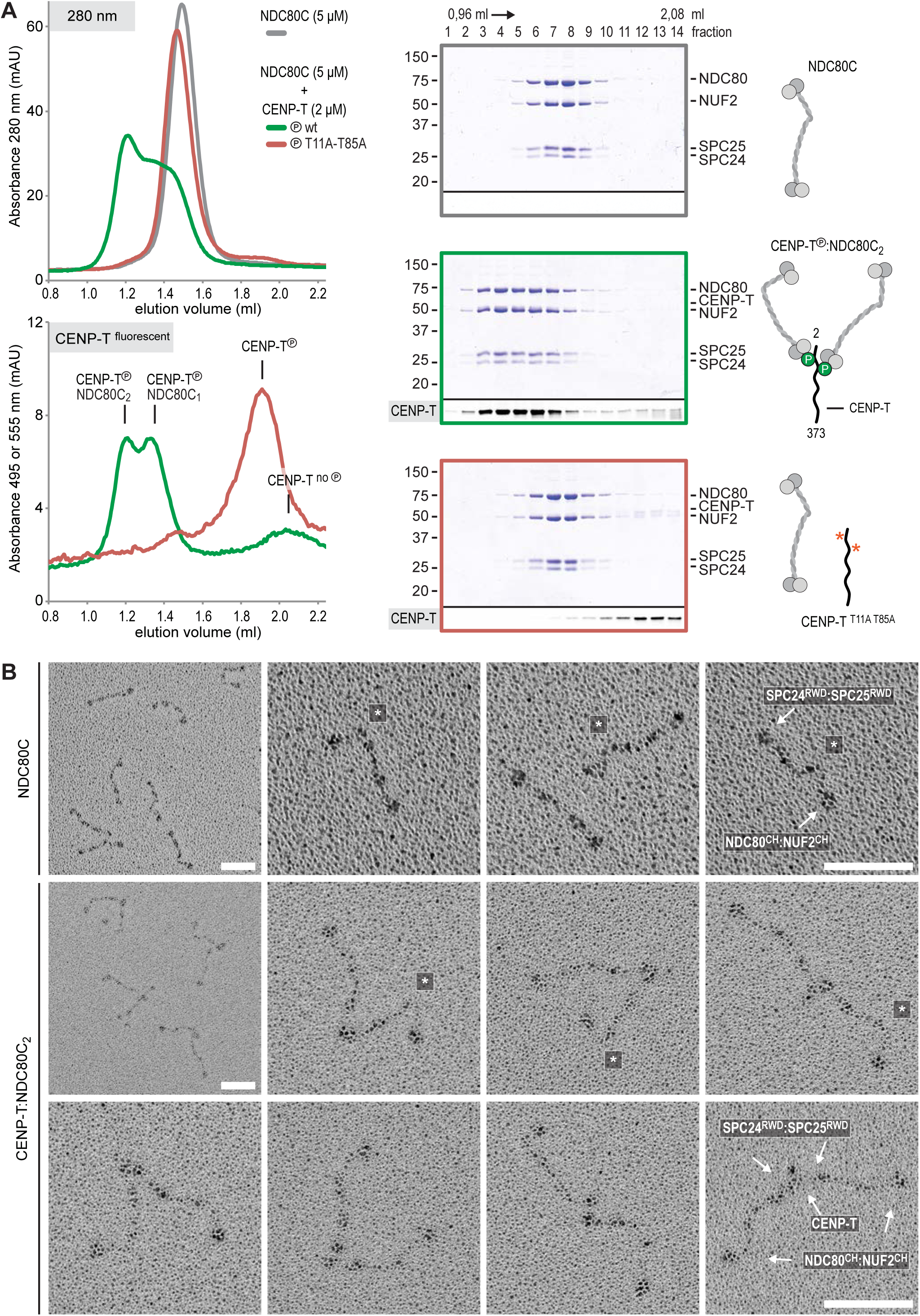
Phosphorylated CENP-T recruits two full-length NDC80 complexes. (**A**). Analytical size-exclusion chromatography (Superose 6 5/150) and SDS-PAGE show that CENP-T^2-373^ phosphorylated by CDK1:Cyclin B at positions T11 and T85 can bind two full-length NDC80 complexes. The mixture of CENP-T (2 µM) and NDC80C (5 µM) contained NDC80C, CENP-T:NDC80C, as well as CENP-T:NDC80C_2_ species (green trace). The specific monitoring of fluorescently labeled CENP-T was used to distinguish these species. Red asterisks indicate mutated phosphorylation sites. Analysis of the single T11A and T85A CENP-T mutants and additional controls are included in **Figure 3 - figure supplement 1**. (**B**) NDC80C (top row) and CENP-T:NDC80C_2_. (middle and bottom rows) were visualized by electron microscopy after glycerol spraying and low-angle platinum shadowing. Asterisks mark the kink in the NDC80:NUF2 coiled coil region. The first micrographs show a representative field of view at a lower magnification. All scale bars represent 50 nm. More micrographs of NDC80C and CENP-T:NDC80C_2_ as well as sample preparation information are included in **Figure 3 - figure supplements 2-4**.

Metal-shadowed particles from an early eluting size exclusion chromatography (SEC) fraction of the CENP-T:NDC80C mixture had a more heterogeneous appearance than NDC80C in isolation (**Figure 3B** and **Figure 3 - figure supplements 3 and 4**). We were able to distinguish at least two different kinds of complexes, containing either one or two NDC80Cs. Though it is not possible to ascertain if complexes with one discernible NDC80 contain CENP-T, those containing two NDC80Cs are presumably assembled on phosphorylated CENP-T. This interpretation is further supported by the distance between the base of the complexes and the characteristic kinks in the coiled coil of NDC80:NUF2 (**Figure 3B** and **Figure 3 - figure supplement 4**). The apparent orientation of the two CENP-T-bound NDC80Cs widely varies, presumably because the CENP-T^11-85^ stretch between the two NDC80C binding sites is flexible. In brief, we visualized how CENP-T recruits two NDC80Cs. The SPC24:SPC25 modules of these complexes are coordinated by the relative proximity of the phosphorylated T11 and T85 of CENP-T, but the microtubule-binding CH domains can be well over 100 nm apart.

### CENP-T phosphorylated at S201 stoichiometrically binds the MIS12 complex

It had previously been shown that MIS12C and CENP-T bind SPC24:SPC25 in a competitive manner (Nishino et al., 2013; Schleiffer et al., 2012). Consistent with these previous reports, we find that MIS12C binds SPC24:SPC25, displacing CENP-T^2-101^, already at equimolar concentrations, indicative of higher binding affinity (unpublished data). However, a similar experiment with the longer CENP-T^2-373^ fragment resulted in binding between MIS12C and phosphorylated CENP-T. This interaction also occurs in the absence of SPC24:SPC25 and strictly depends on the phosphorylation of CENP-T (**Figure 4A** and **Figure 4 - figure supplement 1**, green vs. black traces). Systematic mutational analysis of CENP-T phosphorylation sites (**Table 1**) revealed that the CENPT:MIS12C interaction strictly depends on the phosphorylation of residue S201 of CENP-T (unpublished data and **Figure 4A**, green vs. blue traces). Mutation of the phospho-targets T11 and T85 did not affect the binding between CENP-T and MIS12C (**Figure 4A**, red trace).

**Figure 4.**
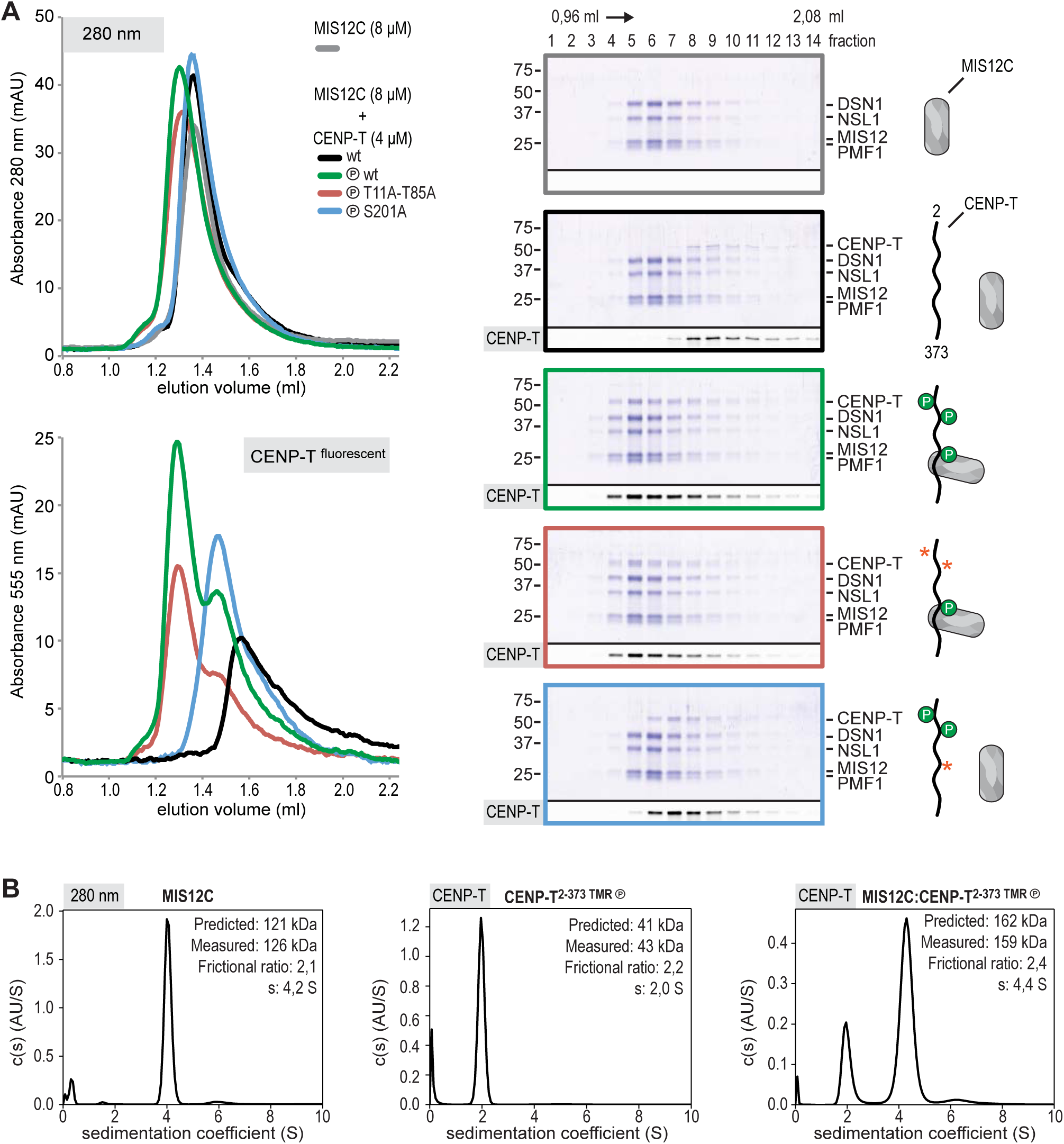
CENP-T phosphorylated by CDK1 at position S201 binds the MIS12 complex. (**A**) Analytical size-exclusion chromatography (Superdex 200 5/150 increase) and SDS-PAGE show that CENP-T^2-373^ phosphorylated by CDK1:Cyclin B at S201 binds to MIS12C. Red asterisks indicate mutated phosphorylation sites. Fluorescently labeled CENP-T was monitored specifically during chromatography and after SDS-PAGE. Additional controls are included in **Figure 4 - figure supplement 1**. (**B**) Sedimentation velocity analytical ultracentrifugation of MIS12C, CENP-T, and MIS12C:CENP-T show that MIS12C and CENP-T bind in a 1:1 complex. MIS12C was monitored by its absorbance at 280 nm whereas CENP-T and the CENP-T:MIS12C mixture were followed using the absorbance of the fluorescently labeled CENP-T^TMR^ at 555 nm.

CENP-T S201 has previously been identified as a mitotic CENP-T phosphorylation site (Kettenbach et al., 2011) and was effectively phosphorylated by CDK1:Cyclin B *in vitro* (**Table 1**), but is not a canonical CDK target site. Notably, phosphorylation of the proximal Thr195, a canonical CDK site that is also phosphorylated *in vitro* (**Table 1**) and *in vivo* (Gascoigne et al., 2011; Kettenbach et al., 2011) and that has been shown to contribute to MIS12C recruitment to CENP-T *in vivo* (Rago et al., 2015), is not required for the direct interaction between CENP-T and MIS12C *in vitro* (**Figure 4 - figure supplement 2**).

We further characterized the interaction between CENP-T and MIS12C using sedimentation velocity analytical ultracentrifugation (SV-AUC) experiments. Both MIS12C and CENP-T sedimented with relatively high frictional ratios, consistent with their elongated shape and unstructured nature, respectively (**Figure 4B**). To analyze the sedimentation of the mixture between CENP-T and MIS12C, we specifically monitored absorbance of the fluorescently labeled CENP-T and thus analyzed sedimentation of CENP-T:MIS12C without following MIS12C. The CENP-T:MIS12C complex sedimented with a frictional ratio of 2.4S and a determined mass (159 kDa), in close agreement with the theoretical mass of a 1:1 complex (162 kDa, unphosphorylated) (**Figure 4B**).

### CENP-T recruits one MIS12 complex and three NDC80 complexes

To test directly if MIS12C that is bound to phosphorylated CENP-T retains its ability to bind NDC80C, we incubated different forms of CENP-T with MIS12C and a molar excess of NDC80C. Phosphorylated CENP-T integrated in large complexes that eluted earlier from the SEC column than MIS12:NDC80 complexes. Our set of CENP-T mutants was used to systematically unravel the complex nature of these CENP-T:MIS12C:NDC80C assemblies. In addition to recapitulating observations made for CENP-T:NDC80C (**Figure 3**) and CENP-T:MIS12C (**Figure 4**), this experiment showed that CENP-T with mutated NDC80C-binding motifs (T11A and T85A) still associated with MIS12C:NDC80C (**Figure 5A**, red trace). This association strictly depends on the phosphorylation of CENP-T at Ser201 because the CENP-T triple mutant (T11A, T85A, S201A) no longer formed a complex with MIS12C or NDC80C (**Figure 5A**, purple trace). Our analyses indicate the formation of assemblies in which CENP-T, MIS12C, and NDC80C are present in 1:1:3 (CENP-T^wt^), 1:1:2 (CENP-T^T11A^ and CENP-T^T85A^), and 1:1:1 (CENP-T^T11A-T85A^) ratios. The behavior of the complete set of CENP-T mutants upon incubation with MIS12C and/or NDC80C is displayed in **Figure 5 - figure supplement 1**).

**Figure 5.**
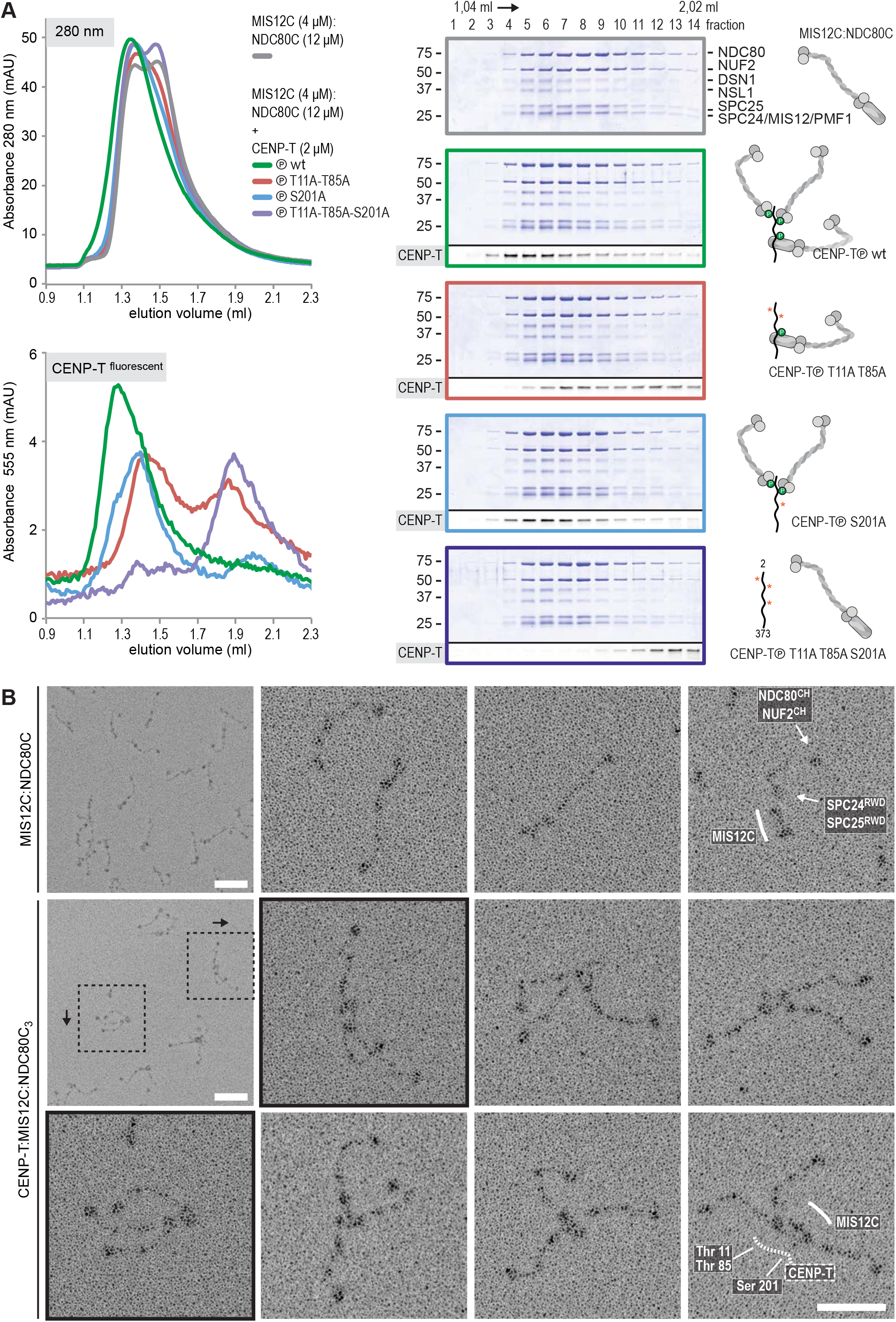
Reconstitution and visualization of CENP-T:MIS12C bound to three full-length NDC80 complexes. (**A**). Analytical size-exclusion chromatography (Superose 6 5/150 increase) and SDS-PAGE show that CENP-T^2-373^ phosphorylated by CDK1:Cyclin B can bind MIS12C and three full-length NDC80 complexes. The triple mutation of phospho-sites T11, T85, and S201 is sufficient to prevent CENP-T from interacting with NDC80C and MIS12C:NDC80C (purple). Red asterisks indicate mutated phosphorylation sites. Fluorescently labeled CENP-T was monitored specifically during chromatography and after SDS-PAGE. The analysis of the complete set of CENP-T mutants is included as **Figure 5 - figure supplement 1**. (**B**) MIS12C:NDC80C (top row) and CENP-T:MIS12C:NDC80C_3_ (middle and bottom rows) were visualized by electron microscopy after glycerol spraying and low-angle platinum shadowing. MIS12C forms a rod-like extension in the MIS12C:NDC80C complexes and can in some cases also be distinguished as a module in CENP-T:MIS12C:NDC80C_3_ assemblies. In those cases, CENP-T T11 and T85 are positioned at the base of the other two NDC80Cs (as annotated in the bottom right micrograph). The first micrographs show a representative field of view at a lower magnification. Scale bars represent 50 nm. Sample preparation is described in **Figure 3 - figure supplement 3**. More micrographs of MIS12C:NDC80C and CENP-T:MIS12C:NDC80C_3_ are included in **Figure 5 - figure supplements 2-3**.

### Visualization of the outer kinetochore assembled on CENP-T

To analyze the overall appearance of CENP-T:MIS12C:NDC80C complexes by electron microscopy, we incubated isolated CENP-T:MIS12C complexes with a molar excess of NDC80C. The mixture was separated by SEC (**Figure 3 - figure supplement 3**) and both a high-molecular weight fraction and a later-eluting fraction were inspected by EM after low-angle metal shadowing. As previously described for complexes that were analyzed by electron microscopy after negative staining (Petrovic et al., 2014; Petrovic et al., 2010; Screpanti et al., 2011), analysis of metal-shadowed MIS12C:NDC80C showed how MIS12C forms a rod-like 18 nm extension at the SPC24:SPC25 side of NDC80C (**Figure 5B** and **Figure 5 - figure supplement 2**). Consistent with our biochemical analysis, the direct visualization of CENP-T:MIS12C:NDC80C assemblies highlighted how CENP-T can coordinate MIS12C and a total of three NDC80Cs into a single complex (**Figure 5B** and **Figure 5 - figure supplement 3**). In a number of cases, the MIS12C:NDC80C module could be identified within the CENP-T:MIS12C:NDC80_3_ assemblies. This enabled the distinction between NDC80C bound to CENP-T directly and NDC80C recruited via the CENP-T:MIS12C interaction. In those cases, the two NDC80Cs that are recruited to CENP-T phosphorylated at positions T11 and T85 resemble the previously visualized CENP-T:NDC80C_2_ complexes (**Figure 5B**, e.g. bottom right).

In a small number of cases, the core of MIS12C:CENP-T seems to be more complex, resulting in assemblies with more density in the middle and four, five, or six NDC80Cs at the periphery (**Figure 5 - figure supplement 3**). Whether this oligomerization represents a biologically relevant tendency of the outer kinetochore to multimerize remains to be tested. Consistent with the aforementioned sample heterogeneity, we also observed CENP-T:MIS12C:NDC80_2_ assemblies (**Figure 5 - figure supplement 3**). These assemblies show one NDC80C that is directly recruited to phosphorylated CENP-T via Thr11 or Thr85 and one NDC80C that binds CENPT:MIS12C. Assemblies with three NDC80Cs were also rare in the later eluting fraction of the same SEC run (**Figure 3 - figure supplement 3**).

### CENP-T and CENP-C are competitive binders of the MIS12 complex

Our observation that CENP-T and MIS12C directly interact raises the possibility that the CENP-C- and CENP-T-mediated outer kinetochore recruitment pathways are integrated into a single CDK1:Cyclin B regulated pathway. However, when we incubated a preformed CENP-T:MIS12C complex with recombinant CENP-C, CENP-T was displaced from MIS12C and a CENP-C:MIS12C complex was formed (**Figure 6** and **Figure 6 - figure supplement 1**). This displacement strictly depended on the ability of CENP-C to bind MIS12C since the CENP-C^K10A-Y13A^ mutant that is unable to bind MIS12C (Screpanti et al., 2011) did not disrupt the CENP-T:MIS12C complex (**Figure 6**) whereas a shorter fragment of CENP-C, CENP-C^1-71^, that can bind MIS12C did (**Figure 6 - figure supplement 1**). Our *in vitro* studies thus demonstrate that MIS12C can be recruited to the kinetochore via CENP-C and via CENP-T that is phosphorylated at position S201. The competitive mode of binding suggests that CENP-C and CENP-T bind to the same site, or at least to partly overlapping sites, of MIS12C, in the latter case in a phosphorylation dependent manner.

**Figure 6.**
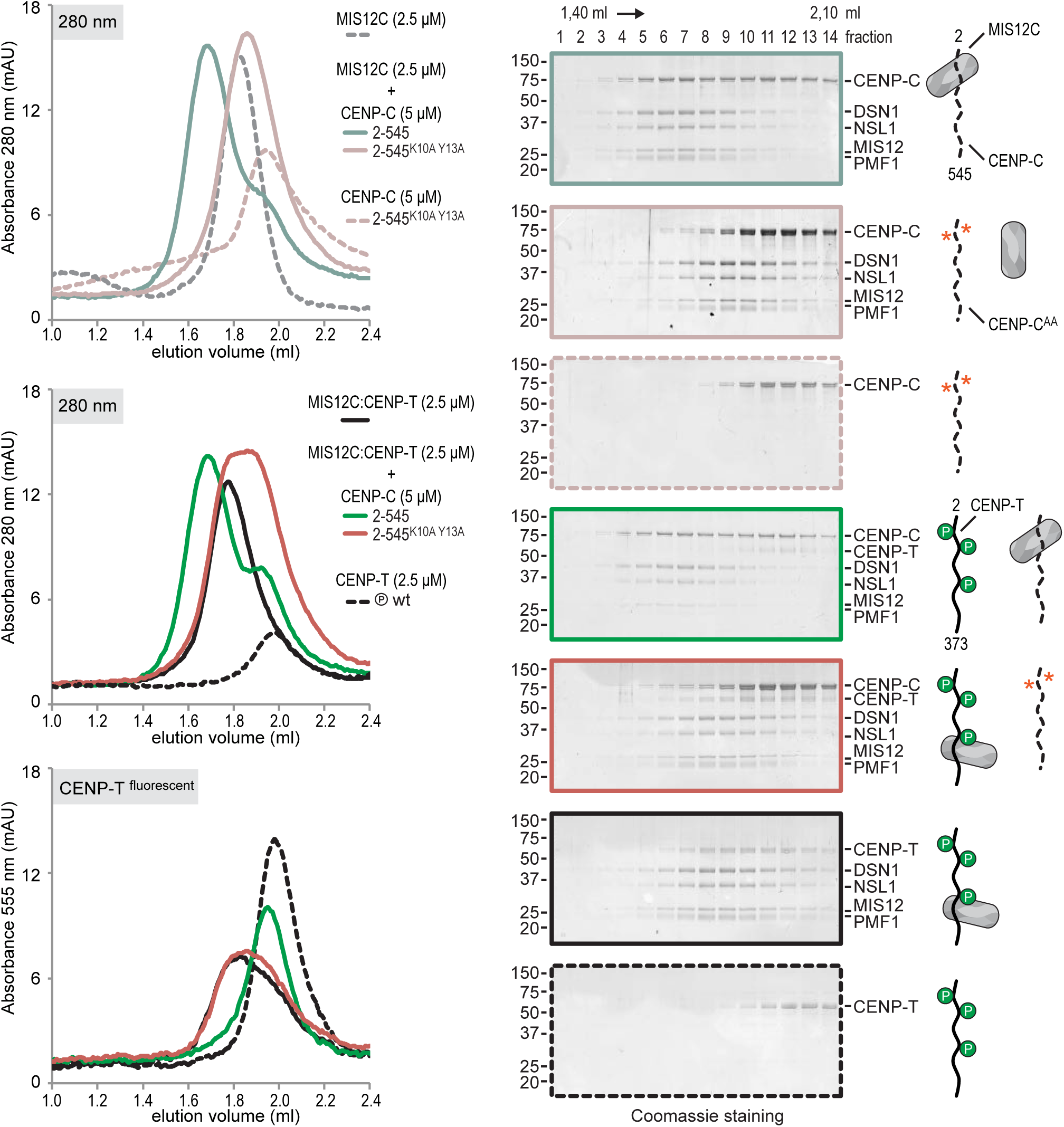
CENP-T and CENP-C are competitive binders of the MIS12 complex. Analytical size-exclusion chromatography (Superdex 200 5/150) and SDS-PAGE show that CENP-T phosphorylated by CDK1-cyclinB at position S201 can bind MIS12C but not MIS12C:CENP-C. Red asterisks indicate mutated phosphorylation sites for CENP-T and the K10A and Y13A mutations in CENP-C. Fluorescently labeled CENP-T was monitored specifically during chromatography.

## Discussion

Full assembly of the outer kinetochore is restricted to mitosis and is tightly regulated, but the molecular requirements for this regulation remain poorly understood. Here, we have reported a thorough dissection of the role of CENP-T in this process. Depletion of CENP-T almost completely abolishes the ultrastructure of the outer kinetochore (Hori et al., 2008) and phenocopies the effects of NDC80C depletion (DeLuca et al., 2005), showing the importance of CENP-T mediated recruitment of NDC80C to mitotic kinetochores. This role is further illustrated by the observation that CENP-T depletion lowers NDC80C levels at mitotic kinetochores more than CENP-C depletion (Gascoigne et al., 2011; Suzuki et al., 2015), and by experiments showing that replacement of the outer kinetochore binding domain of CENP-C with that of CENP-T supports chromosome segregation in cells depleted of CENP-C (Suzuki et al., 2015).

We reconstituted the assembly of outer kinetochore modules using purified components and determined how CDK-dependent phosphorylation of CENP-T controls the multivalency of NDC80C. The analysis of our synthetic outer kinetochores by SEC, SV-AUC, and EM revealed that one copy of CENP-T recruits two NDC80Cs via phosphorylated CENP-T^T11^ and CENP-T^T85^ and one MIS12C:NDC80C via phosphorylated CENP-T^S201^. Because the binding of CENP-C and CENP-T to MIS12C is competitive, these findings support a model in which CENP-C and CENP-T act in parallel to recruit two MIS12Cs and up to four NDC80Cs in a CDK-regulated manner (**Figure 7**).

**Figure 7.**
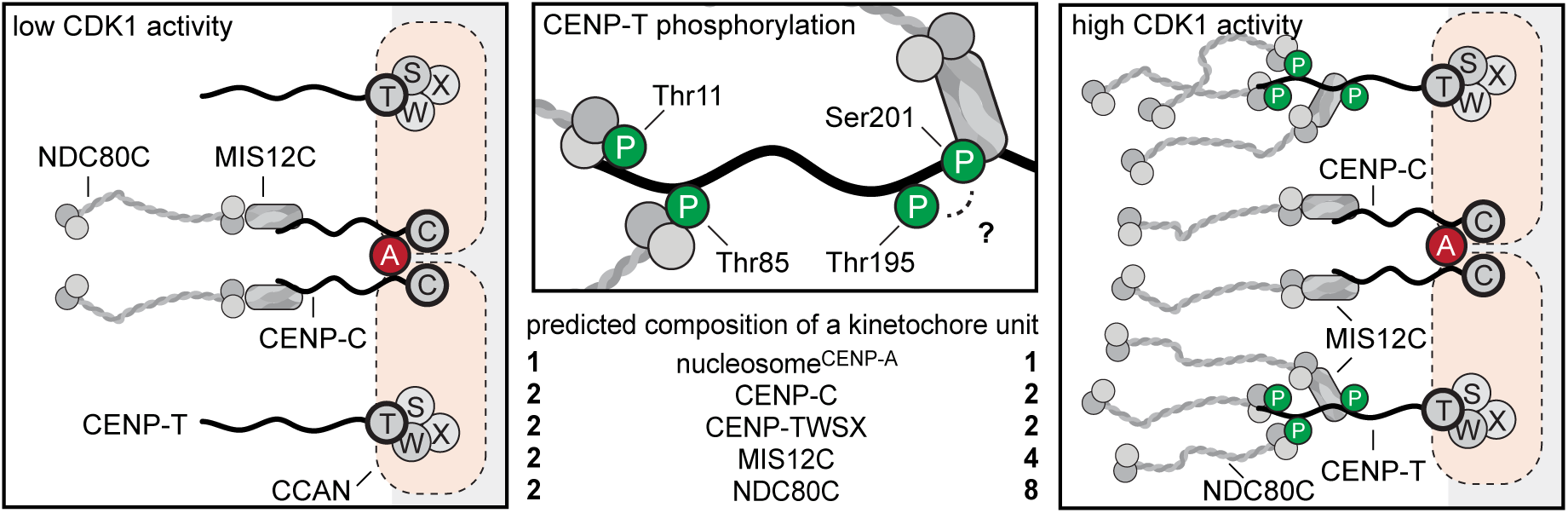
Molecular basis of stoichiometric assembly of microtubule binders in the outer kinetochore. Schematic representation of the results presented here and their implications. CDK activity results in the recruitment of NDC80C and MIS12C:NDC80C to CENP-T. The predicted stoichiometry of a single kinetochore unit is indicated. In the middle panel, we summarize the role of T11, T85, and S201 in recruitment of NDC80C and MIS12C, and suggest that T195 may contribute to phosphorylation of S201 *in vivo*.

Our finding that phosphorylated CENP-T can form a 1:1 complex with MIS12C is consistent with a series of *in vivo* studies that showed that CENP-T does not only bring NDC80C to the outer kinetochore, but is also involved in the recruitment of MIS12C (Gascoigne and Cheeseman, 2013; Gascoigne et al., 2011; Hori et al., 2013; Kim and Yu, 2015; Rago et al., 2015; Suzuki et al., 2015). The molecular mechanism behind the contribution of CENP-T to MIS12C localization at the kinetochore, however, had remained unclear, and it appeared confusing in view of observations that MIS12C and CENP-T bind to NDC80C in a competitive manner (Nishino et al., 2013; Schleiffer et al., 2012). A role for the CENP-T^200-230^ region in MIS12C recruitment was shown in experiments using the ectopic chromatin targeting of CENP-T fragments that revealed that MIS12C was recruited to CENP-T^1-230^ but not to CENP-T^1-200^ (Rago et al., 2015). Our observation that the phosphorylation of S201 is essential for the CENP-T:MIS12C interaction *in vitro* explains these previous observations (**Figure 4**). Interestingly, phosphorylation of the CDK consensus site CENP-T T195 is important for the CENPT:MIS12C interaction *in vivo*(Rago et al., 2015), but is not required *in vitro* (**Figure 4 - figure supplement 2**). A possible explanation for these results is that the phosphorylation of kinetochore-associated CENP-T at Thr195 is required for the phosphorylation of Ser201 and the subsequent binding of MIS12C. Multisite phosphorylation networks are important for the regulation of mitosis by CDK1 kinase (Koivomagi et al., 2011) and future experiment will be required to understand the spatial and temporal regulation of CENP-T phosphorylation *in vivo*.

Our reconstitution experiments provide a molecular explanation for the observed CDK-dependent recruitment of NDC80C to kinetochores (Gascoigne et al., 2011) and show how one CENP-T molecule can recruit one MIS12C and up to three NDC80Cs. Because the pseudo-symmetric nature of CENP-A nucleosomes results in the binding of two CCAN modules (Weir et al., 2016), we propose that mitotic phosphorylation events trigger the recruitment of four MIS12Cs and up to eight NDC80Cs per CENP-A mononucleosome (**Figure 7**). This extrapolation of our reconstitution experiments is in line with a series of quantitative analyses of kinetochore subcomplexes by immunofluorescence microscopy. Copy numbers of CENP-C, CENP-T, MIS12C, and NDC80C were most accurately determined for metaphase kinetochores in a study that combined RNA interference, ectopic and endogenous kinetochore recruitment, and an elegant CENP-C/T chimera protein design (Suzuki et al., 2015). This analysis revealed that, on average, 215 copies of CENP-C and 72 copies of CENP-T recruit a total of 244 NDC80Cs and 151 MIS12Cs to a kinetochore. However, only 82 (of the 215) CENP-C proteins engage in binding MIS12C, whereas the remaining CENP-C does not contribute to outer kinetochore recruitment. CENP-T is responsible for the recruitment of the remaining MIS12Cs and NDC80Cs (Suzuki et al., 2015). These results correlate well to our description of 1:1:1 CENP-C:MIS12C:NDC80C and 1:1:2 or 1:1:3 CENP-T:MIS12C:NDC80C assemblies.

As stated in the introduction, NDC80C multivalency may have profound implications for microtubule binding (Hill, 1985; Zaytsev et al., 2014). Our study paves the way to the reconstitution of synthetic kinetochores with controllable numbers of NDC80 complexes, which will in turn allow us to probe the importance of multivalency for force production by kinetochores. Whether the CDK-regulated NDC80C multivalency at the outer kinetochore has an effect on the function of kinetochores to signal the microtubule-attachment state through spindle checkpoint control also remains to be addressed. In this perspective, it is interesting to note that NDC80C recruited by CENP-C:MIS12C or by CENP-T have been proposed to have different effects on the checkpoint response (Kim and Yu, 2015; Samejima et al., 2015).

The visualization of the reconstituted 1:1:3 CENP-T:MIS12C:NDC80C complexes by electron microscopy after low-angle metal shadowing highlighted how outer kinetochores assembled on phosphorylated CENP-T can span distances of well over 100 nm. It also showed the varying orientation of the coordinated NDC80Cs relative to each other, presumably caused by the flexibility of the CENP-T regions in between the NDC80C and MIS12C binding sites. The flexibility of CENP-T as well as the stretching of CENP-T upon kinetochore-microtubule contacts has been shown before: the N- and C-termini of tagged CENP-T were ~30 nm apart in the presence of tension (MG132 treated cells) but only ~4 nm apart in the absence of tension (nocodazole treated cells), as determined by immunofluorescence and immuno-electron microscopy (Suzuki et al., 2011). Moreover, the N-terminal tail of CENP-T was seen as a 25 ± 13 nm long flexible extension of the CENP-TW histone-fold domain by high speed atomic force microscopy (Suzuki et al., 2011).

The likely interpretation of these results is that the intrinsically flexible N-terminal region of CENP-T stretches when the multiple NDC80Cs it coordinates bind microtubules, while its C-terminal HFD remains anchored at the inner kinetochore. In the future, it will be important to test how the observed CENP-T:MIS12C:NDC80C_3_ complexes structurally rearrange in the presence of microtubules and if the presence of multiple NDC80Cs contributes to stabilization of the kinetochore-microtubule interface observed in the presence of tension (Akiyoshi et al., 2010; Nicklas and Koch, 1969). Low affinity, cooperative interactions might acquire the highest relevance under these conditions, and biochemical reconstitution is a powerful means to identify them and address their possible importance.

## Experimental Procedures

### Protein expression and purification

cDNAs encoding GST-CENP-T^2-101^ and GST-CENP-T^2-373^ with a C-terminal -LPETGG extension were subcloned into pGEX-6P-2rbs, a dicistronic derivative of pGEX6P, generated in house. Point mutations were introduced by PCR and all plasmids were verified by DNA sequencing. *E. coli* BL21(DE3)-Codon-plus-RIL cells containing the CENP-T encoding pGEX-6P-2rbs vector were grown at 37°C in Terrific Broth in the presence of Chloramphenicol and Ampicillin to an OD_600_ of ~0.8. Protein expression was induced by the addition of 0.35 mM IPTG and cells were incubated ~14 hours at 20°C. All following steps were performed on ice or at 4°C. Cell pellets were resuspended in buffer A (20 mM Tris-HCl, pH 8.0, 300 mM NaCl, 10% (v/v) glycerol and 1 mM TCEP) supplemented with 0.5 mM PMSF, protease-inhibitor mix HP Plus (Serva) and DNaseI (Roche), lysed by sonication and cleared by centrifugation at 108,000*g* for 30 minutes. The cleared lysate was applied to a 5 ml GSTrap FF column (GE Healthcare) equilibrated in buffer A. The column was washed with approximately 50 column volumes of buffer A and bound protein was eluted in buffer A by cleavage of the GST-tag with PreScission protease for ~14 hours. The eluate was diluted 10-fold with buffer B (20 mM Tris-HCl pH 6.8, 1 mM TCEP) and applied to a 5 ml HiTrap Heparin HP column (GE Healthcare) pre-equilibrated in buffer B with 25 mM NaCl. Bound protein was eluted in buffer B using a linear gradient from 25 mM to 800 mM NaCl in 16 column volumes. Selected fractions were concentrated (Amicon; 10 kDa molecular weight cut-off) and either used for Sortase-mediated labeling (see below) or directly applied to an equilibrated Superdex 200 10/300 SEC column (GE Healthcare). SEC was performed with a 20 mM Tris-HCl, pH 8.0, 150 mM NaCl, and 1 mM TCEP mobile phase at a flow rate of 0.4 ml/min, and the relevant fractions were pooled, concentrated, flash-frozen in liquid nitrogen, and stored at -80 °C.

The four full-length components of the NDC80 complex (NDC80C; SPC25^6xHis^) were combined on a pFL (Fitzgerald et al., 2006) or a pBIG1 (Weissmann et al., 2016) vector. Baculoviruses were generated in Sf9 insect cells and used for protein expression in Tnao38 insect cells (Hashimoto et al., 2010). Between 60 and 72 hours post-infection, cells were pelleted, washed in PBS, pelleted, and stored at -80 °C until use. Cells were thawed and resuspended in buffer A (50 mM Hepes, pH 8.0, 200 mM NaCl, 5% v/v glycerol, 1 mM TCEP) supplemented with 20 mM imidazole, 0.5 mM PMSF, and protease-inhibitor mix HP Plus (Serva), lysed by sonication and cleared by centrifugation at 108,000*g* for 30 minutes. The cleared lysate was filtered (0.8 µM) and applied to a 5 ml HisTrap FF (GE Healthcare) equilibrated in buffer A with 20 mM imidazole. The column was washed with approximately 50 column volumes of buffer A with 20 mM imidazole and bound proteins were eluted in buffer A with 300 mM imidazole. Relevant fractions were pooled, diluted 5-fold with buffer A with 25 mM NaCl and applied to a 6 ml ResourceQ column (GE Healthcare) equilibrated in the same buffer. Elution of bound protein was achieved by a linear gradient from 25 mM to 400 mM NaCl in 30 column volumes. Relevant fractions were concentrated in 30 kDa molecular mass cut-off Amicon concentrators (Millipore) in the presence of an additional 200 mM NaCl and applied to a Superdex 200 10/300 or a Superose 6 10/300 column (GE Healthcare) equilibrated in size-exclusion chromatography buffer (50 mM Hepes, pH 8.0, 250 mM NaCl, 5% v/v glycerol, 1 mM TCEP). Size-exclusion chromatography was performed under isocratic conditions at recommended flow rates and the relevant fraction were pooled, concentrated, flash-frozen in liquid nitrogen, and stored at -80 °C. Expression and purification of the full-length MIS12 complex (MIS12C; Dsn1^6His^) was identical but size exclusion chromatography was performed in 50 mM Hepes, pH 8.0, 200 mM NaCl, 1 mM and no additional NaCl was added before the concentration of the relevant fractions of the ion-exchange chromatography step.

SPC24:SPC25 (Ciferri et al., 2008), CENP-C1^-71^ (Screpanti et al., 2011) and CENP-C^2-545^ (Klare et al., 2015) were expressed and purified as described.

Codon-optimized CDK1 and Cyclin B1 constructs were obtained from GeneArt (Life Technologies) and combined in a pBIG1A vector (Weissmann et al., 2016) with N-terminal tags on CDK1 (GST) and Cyclin B1 (hexahistidine). GST-CDK1 and 6His-Cyclin B1 were co-expressed in Tnao38 insect cells as described above and purified using glutathione sepharose (GE Healthcare) followed by size exclusion chromatography using a HiLoad 16/60 Superdex 200 pg column (GE Healthcare) equilibrated in buffer containing 20 mM HEPES pH 7.5, 200 mM NaCl, 1 mM TCEP, and 5% glycerol.

### Sortase labeling

Purified *S. aureus* Sortase (Guimaraes et al., 2013) or the Sortase 5M mutant (Hirakawa et al., 2015) were used to label CENP-T^2-373^-LPETGG with GGGGK peptides with a C-terminally conjugated tetramethylrhodamine (TMR) or fluorescein amidite (FAM) (Genscript). Labeling was performed for ~14 hours at 4°C in the presence of 10 mM CaCl _2_using molar ratios of Sortase, CENP-T, and peptide of approximately 1:20:200. CENP-T was separated from Sortase and the unreacted peptides by size exclusion chromatography as described above. Based on the absorbance at 280 nm and 495 nm (FAM) or 555 nm (TMR), a labeling efficiency of >90% was achieved. Sortase labeling of SPC24:SPC25 was performed in a similar manner.

### *In vitro* phosphorylation of CENP-T

CENP-T^2-101^ fragments and CENP-T analyzed by mass spectrometry were *in vitro* phosphorylated by CDK1:Cyclin B (Millipore). In-house generated CDK1:CyclinB was used for all other *in vitro* phosphorylation experiments. Phosphorylation reactions were set up in size exclusion chromatography buffer (20 mM Tris-HCl pH 8, 150 mM NaCl, and 1 mM TCEP) containing 100 nM CDK1:Cyclin B, 10 µM CENP-T substrate, 2 mM ATP, and 10 mM MgCl_2_. Reaction mixtures were kept at 30°C for 30 minutes and then used in binding experiments.

### Mass spectrometry

Liquid chromatography coupled with mass spectrometry was used to asses the phosphorylation status of *in vitro* phosphorylated CENP-T^2-373^. An unposphorylated sample was processed in parallel and used as a control. Samples were reduced, alkylated and digested with GluC or LysC/Trypsin and prepared for mass spectrometry as previously described (Rappsilber et al., 2007). Obtained peptides were separated on an EASY-nLC 1000 HPLC system (Thermo Fisher Scientific, Odense, Denmark) using a 45 min gradient from 5-60% acetonitrile with 0.1% formic acid and directly sprayed via a nano-electrospray source in a quadrupole Orbitrap mass spectrometer (Q Exactive, Thermo Fisher Scientific) (Michalski et al., 2011). The Q Exactive was operated in data-dependent mode acquiring one survey scan and subsequently ten MS/MS scans. Resulting raw files were processed with the MaxQuant software (version 1.5.2.18) using CENP-T^2-373^ for the search with deamidation (NQ), oxidation (M) and phosphorylation (STY) as variable modifications and carbamidomethylation (C) as fixed modification. A false discovery rate cut off of 1% was applied at the peptide and protein levels and as well on the site decoy fraction (Cox and Mann, 2008).

### Analytical size exclusion chromatography

Proteins were mixed at the indicated concentrations, incubated on ice for at least one hour, spun for 15 minutes at 13,000 rpm at 4 °C, and then analyzed by size exclusion chromatography at 4 °C using a ÄKTAmicro system (GE Healthcare) mounted with a column (Superdex 200 5/150, Superdex 200 5/150 increase, Superose 6 5/150, or Superose 6 5/150) equilibrated in size exclusion chromatography buffer (20 mM Tris-HCl pH 8, 150 mM NaCl, and 1 mM TCEP) and operated at or near the recommended flow rate. Fractions of 50-100 µL were collected in 96-well plates and analyzed by SDS-PAGE. The absorbance of fluorescently labeled proteins was followed during chromatography using the ÄKTAmicro Monitor UV-900 (GE Healthcare) and after SDS-PAGE using a ChemiDoc MP system (Bio-Rad).

### Isothermal titration calorimetry

All samples were exchanged into fresh buffer (20 mM Tris-HCl, 150 mM NaCl and 1 mM TCEP). ITC measurements were performed at 20 °C on an ITC200 microcalorimeter (GE Healthcare). In each titration, the protein in the cell (at a 10 - 30 µM concentration) was titrated with 25 x 4 µl injections (at 180 sec intervals) of protein ligand (at 10-fold higher molar concentration). The following synthetic peptides (purity > 95 %, Genscript) were used Thr11 DST(p)PRTLLRRVLDTAYA, Thr85 EQT(p)PRTLLKNILLTAYA, Thr27 PRT(p)PRRPRSARAGARYA, and Thr47 TAS(p)PRKLSGQTRTIARYA. Injections were continued beyond saturation to allow for determination of heats of ligand dilution. Data were fitted by least-square procedure to a single-site binding model using ORIGIN software package (MicroCal).

### Analytical Ultracentrifugation

Sedimentation velocity AUC was performed at 42,000 rpm at 20°C in a Beckman XL-A ultracentrifuge. Purified protein samples were diluted to 15-30 µM in a buffer containing 20 mM Tris-HCl, 150 mM NaCl and 1 mM TCEP and loaded into standard double-sector centerpieces. The cells were scanned at 280 nm or 555 nm every minute and 500 scans were recorded for every sample. Data were analyzed using the program SEDFIT (Schuck, 2000) with the model of continuous *c*(*s*) distribution. The partial specific volumes of the proteins, buffer density and buffer viscosity were estimated using the program SEDNTERP. Data figures were generated using the program GUSSI.

### Low-angle metal shadowing and electron microscopy

Fractions from an analytical size exclusion chromatography column were diluted 1:1 with spraying buffer (200 mM ammonium acetate and 60% glycerol) and air-sprayed as described (Baschong and Aebi, 2006) onto freshly cleaved mica (V1 quality, Plano GmbH) of approximately 2x3 mm. Specimens were mounted in a MED020 high-vacuum metal coater (Bal-tec) and dried for ~14 hours. A Platinum layer of approximately 1 nm and a 7 nm Carbon support layer were subsequently evaporated onto the rotating specimen at angles of 6-7° and 45° respectively. Pt/C replicas were released from the mica on water, captured by freshly glow-discharged 400-mesh Pd/Cu grids (Plano GmbH), and visualized using a LaB_6_ equipped JEM-400 transmission electron microscope (JEOL) operated at 120 kV. Images were recorded at a nominal magnification of 60,000x on a 4k X 4k CCD camera F416 (TVIPS), resulting in 0.1890 nm per pixel. Particles were manually selected using EMAN2 (Tang et al., 2007) and further analyzed using Fiji (Schindelin et al., 2012).

## Author contributions

P.J.H., S.J., A.P., J.J., and A.M. designed, performed, and interpreted experiments. S.J., P.J.H., J.J., A.P., and P.S. purified proteins. P.J.H., S.J., and J.J. performed analytical size-exclusion chromatography. P.J.H. performed low-angle metal shadowing and electron microscopy. A.P. and J.J. performed analytical ultracentrifugation. A.P. and S.J. performed isothermal titration calorimetry. F.W. contributed novel reagents. T.B. performed mass spectrometry. P.J.H. and A.M. prepared figures and wrote the manuscript with input from all authors.

## Acknowledgements

We are grateful to S. Wohlgemuth and I. Stender for NDC80C and MIS12C purifications and to O. Hofnagel and F. Müller for assistance with electron microscopy and mass spectrometry. We thank J.-M. Peters for sharing unpublished reagents and acknowledge S. van Gerwen, J. Keller, K. Klare, V. Krenn, A. Maiolica, S. Mosalaganti, A. Schleiffer, and F. Villa, as well as all members of the Musacchio laboratory for support, ideas, and discussion. S.J. acknowledges support of a Marie Curie Mobility Fellowship, SEMM’s Structured International Post Doc program (SIPOD) and an EMBO long-term fellowship (ALTF 262-2009). A.M. acknowledges funding by the European Union’s 7th Framework Program ERC advanced grant agreement RECEPIANCE and the DFG’s Collaborative Research Centre (CRC) 1093. The authors declare no competing financial interests.

---

**Figure 1 - figure supplement 1.**
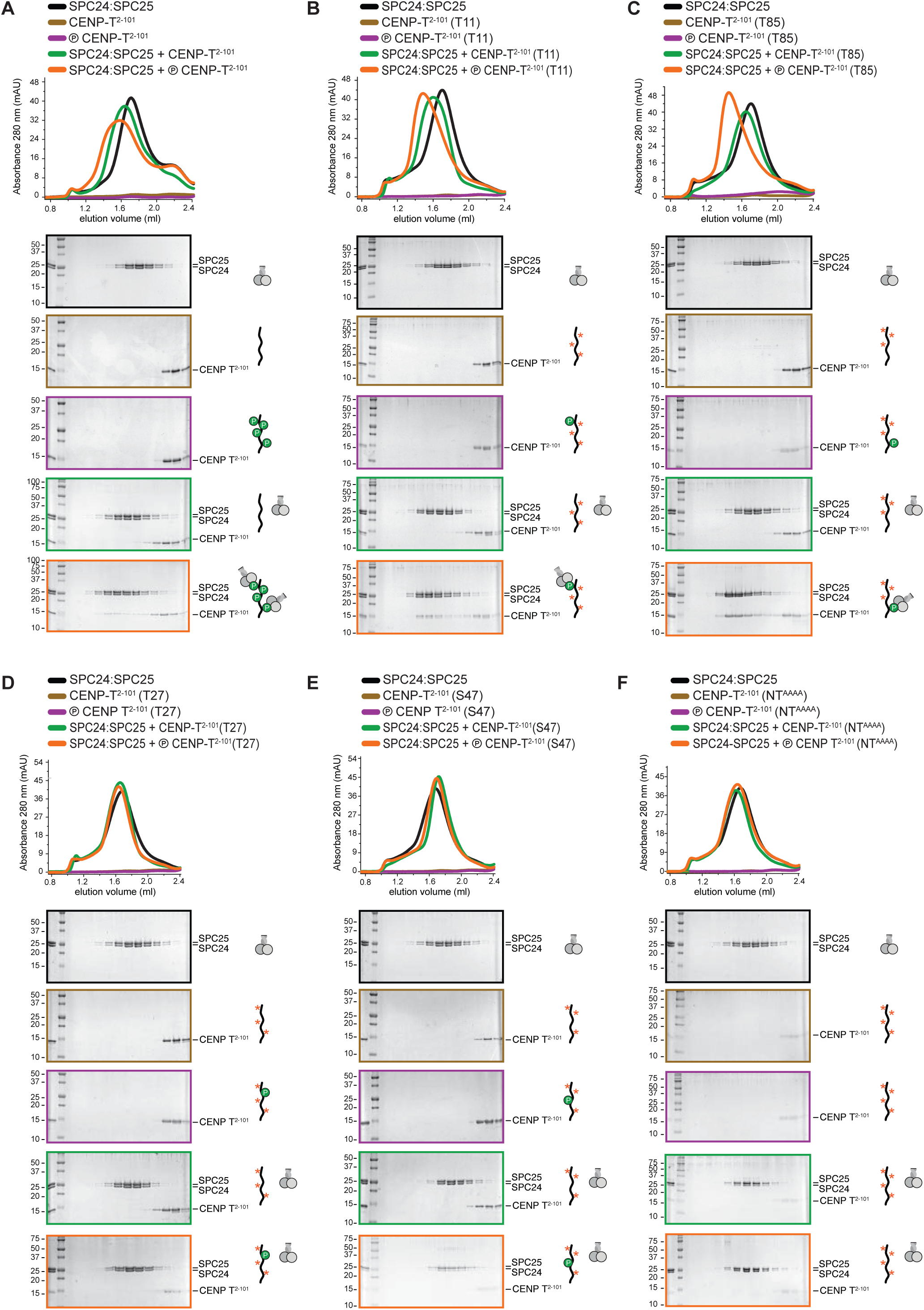
Phosphorylation of CENP-T^2-101^ at T11 or T85 is sufficient for the binding of SPC24:SPC25. SDS-PAGE analysis of various CENP-T^2-101^ mutants that were incubated with SPC24:SPC25 and separated by analytical size-exclusion chromatography (Superdex 200 5/150). The six gels boxed in orange are displayed in Figure 1D.

**Figure 3 - figure supplement 1.**
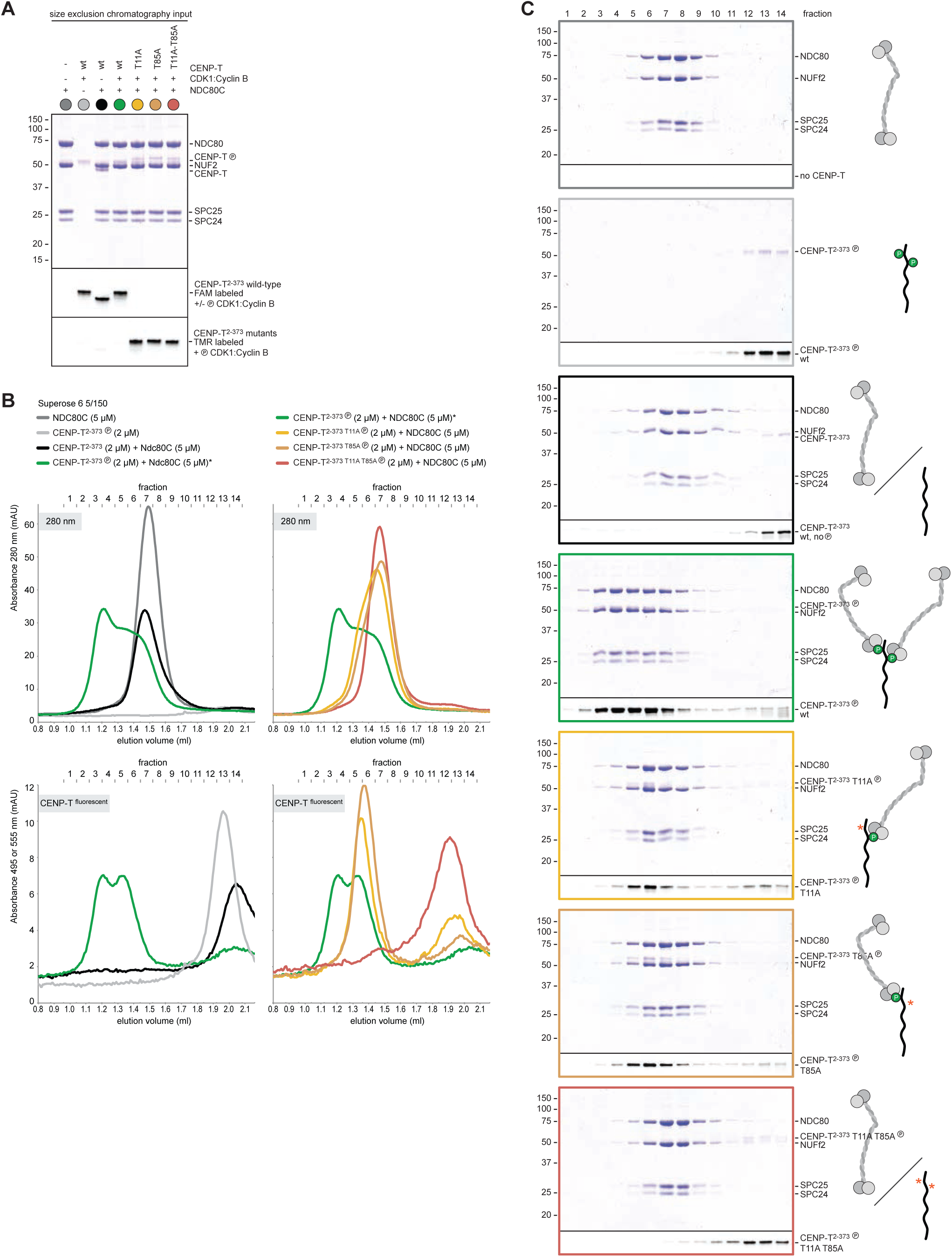
Phosphorylated CENP-T recruits two full-length NDC80 complexes. (**A**) A fraction of the samples was analyzed by SDS-PAGE prior to size exclusion chromatography. The phosphorylation-induced shift of CENP-T is clearly visible. (**B**, **C**) Analytical size-exclusion chromatography (Superose 6 5/150) and SDS-PAGE showing that CENP-T phosphorylated by CDK1 at positions T11 and T85 can bind two full-length NDC80 complexes. Data for NDC80C alone (grey), NDC80C mixed with phosphorylated wild-type (green) or T11A-T85A (red) CENP-T are displayed in **Figure 3**.

**Figure 3 - figure supplement 2.**
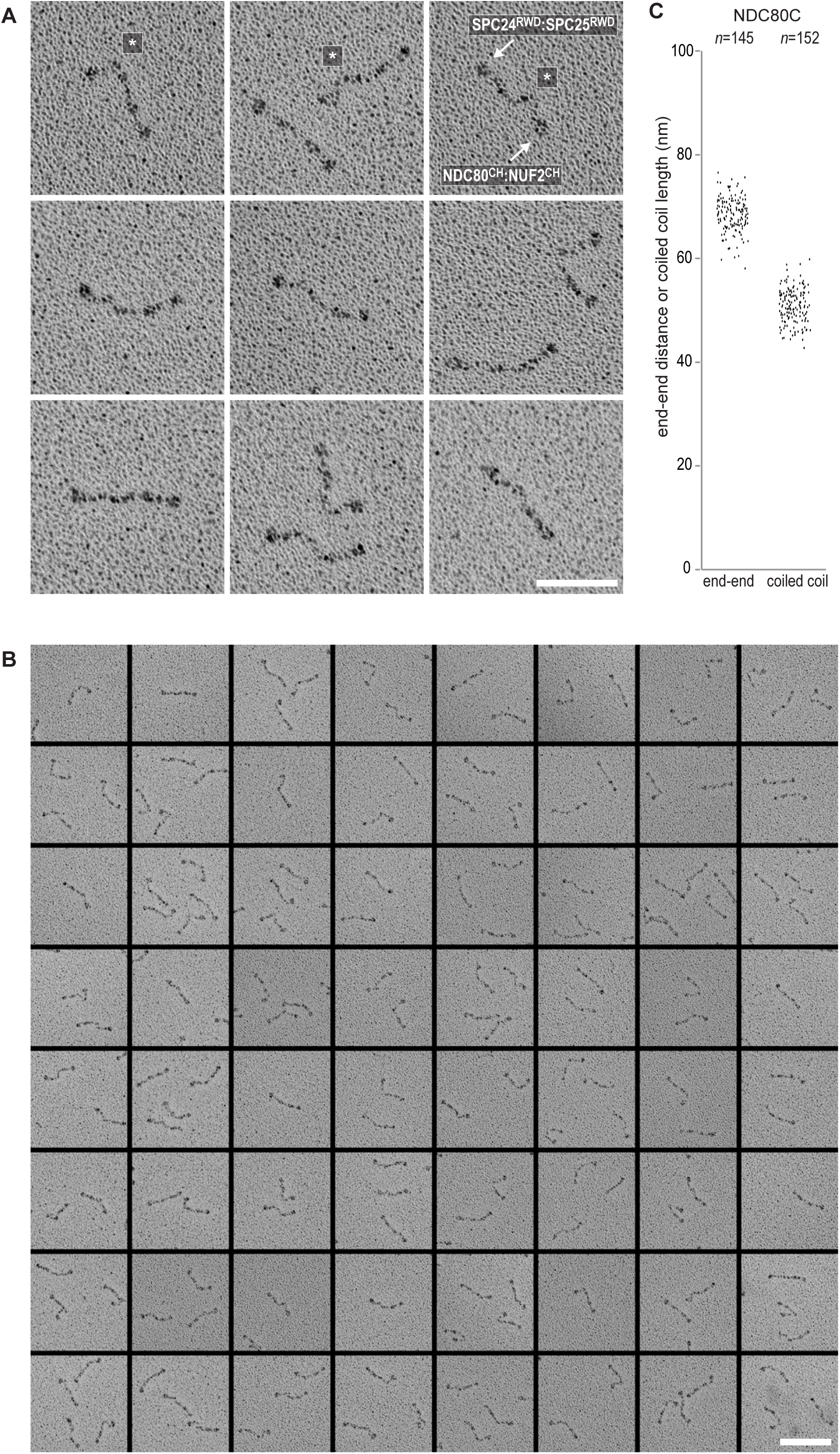
Gallery and measurements of NDC80C. (**A**) Purified NDC80 complexes were separated by size exclusion chromatography and visualized after glycerol spraying followed by low-angle metal shadowing. The three micrographs in the top row are shown in **Figure 3**. (**B**) Gallery showing 64 selected areas with one or more NDC80Cs. Scale bars represent 50 nm. (**C**) End-end and coiled coil distances were measured for the indicated number of shadowed NDC80Cs. The displayed values are not corrected for an estimated low-angle shadowing contribution of 3 nm at either end of NDC80C.

**Figure 3 - figure supplement 3.**
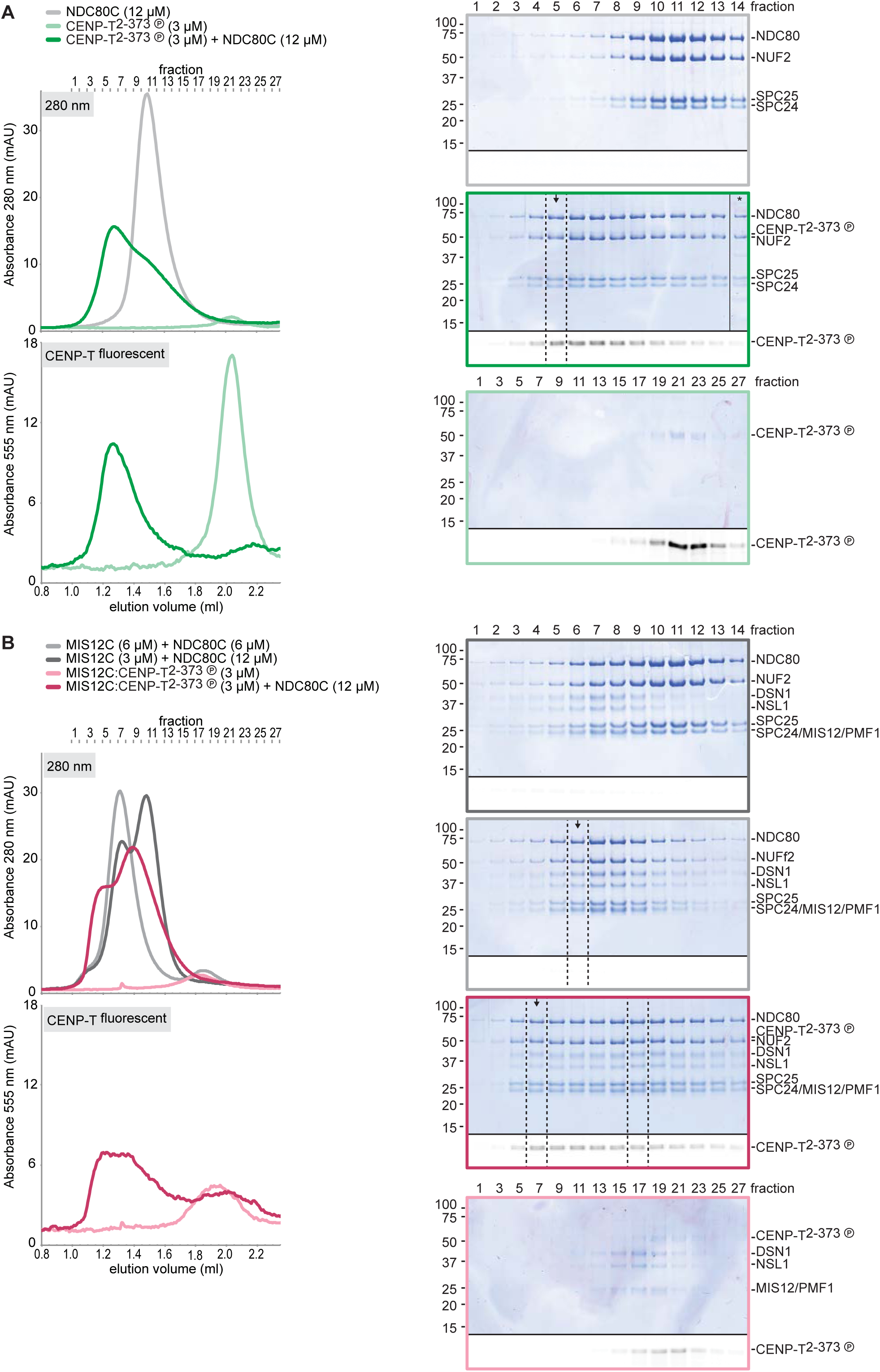
Sample preparation for shadowing EM. NDC80C was mixed with phosphorylated CENP-T or with an isolated CENP-T:MIS12C complex and analyzed by analytical size-exclusion chromatography (Superose 6 5/150) and SDS-PAGE. Different gels from the same material were used for Coomassie staining and the stain-free identification of fluorescent CENP-T. Samples that were used for low-angle metal shadowing are indicated by the black arrow. Results are shown in **Figure 3B** and **Figure 5B** as well as in **Figure 3 - figure supplement 4** and **Figure 5 - figure supplements 2-3**.

**Figure 3 - figure supplement 4.**
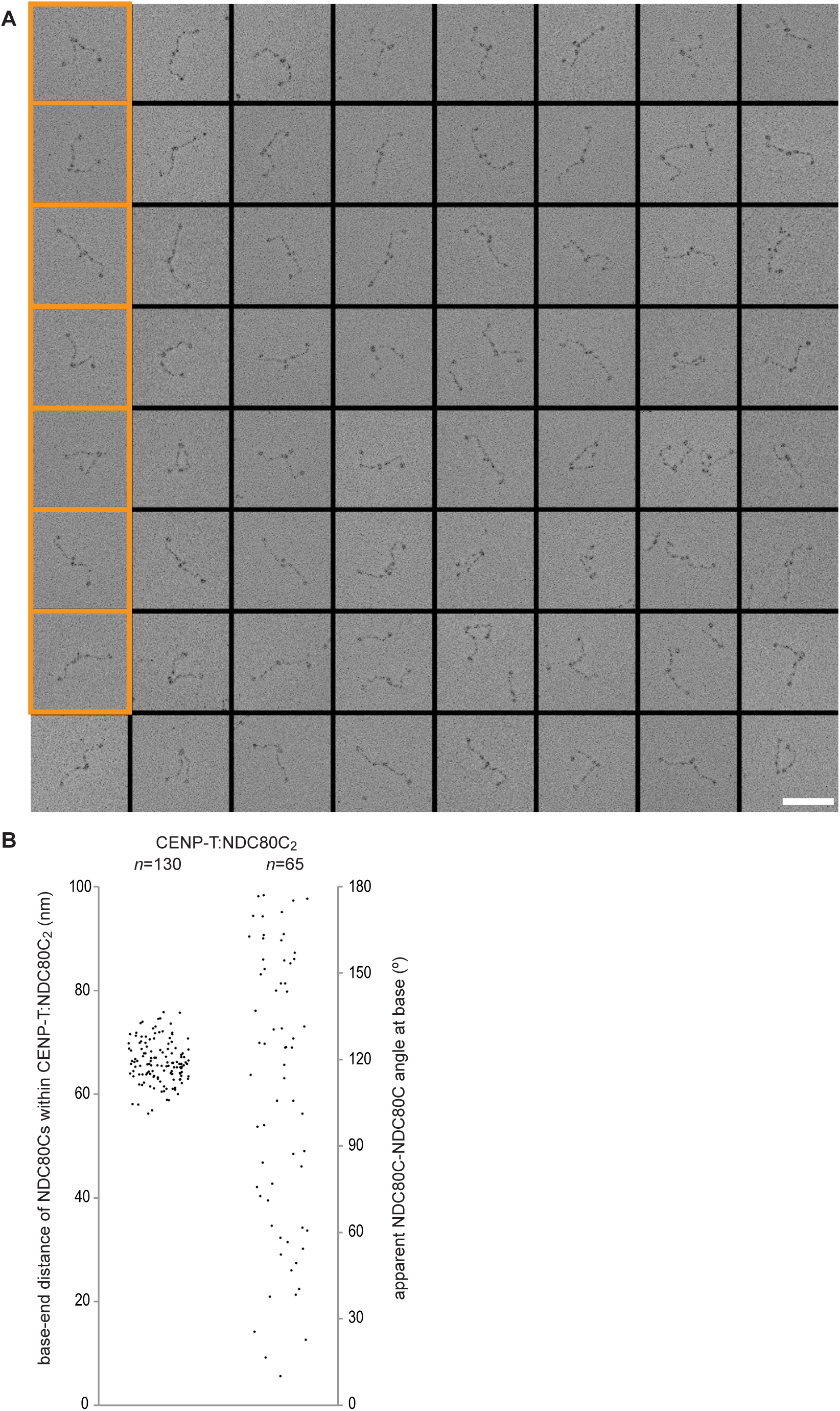
Gallery and measurements of CENP-T:NDC80C_2_. (**A**) CENP-T:NDC80C_2_ complexes were visualized after glycerol spraying followed by low-angle metal shadowing. The orange-boxed micrographs are shown in **Figure 3B**. Scale bar 50 nm. (**B**) The apparent relative orientation of the two NDC80Cs in CENP-T:NDC80C_2_ complexes was measured.

**Figure 4 - figure supplement 1.**
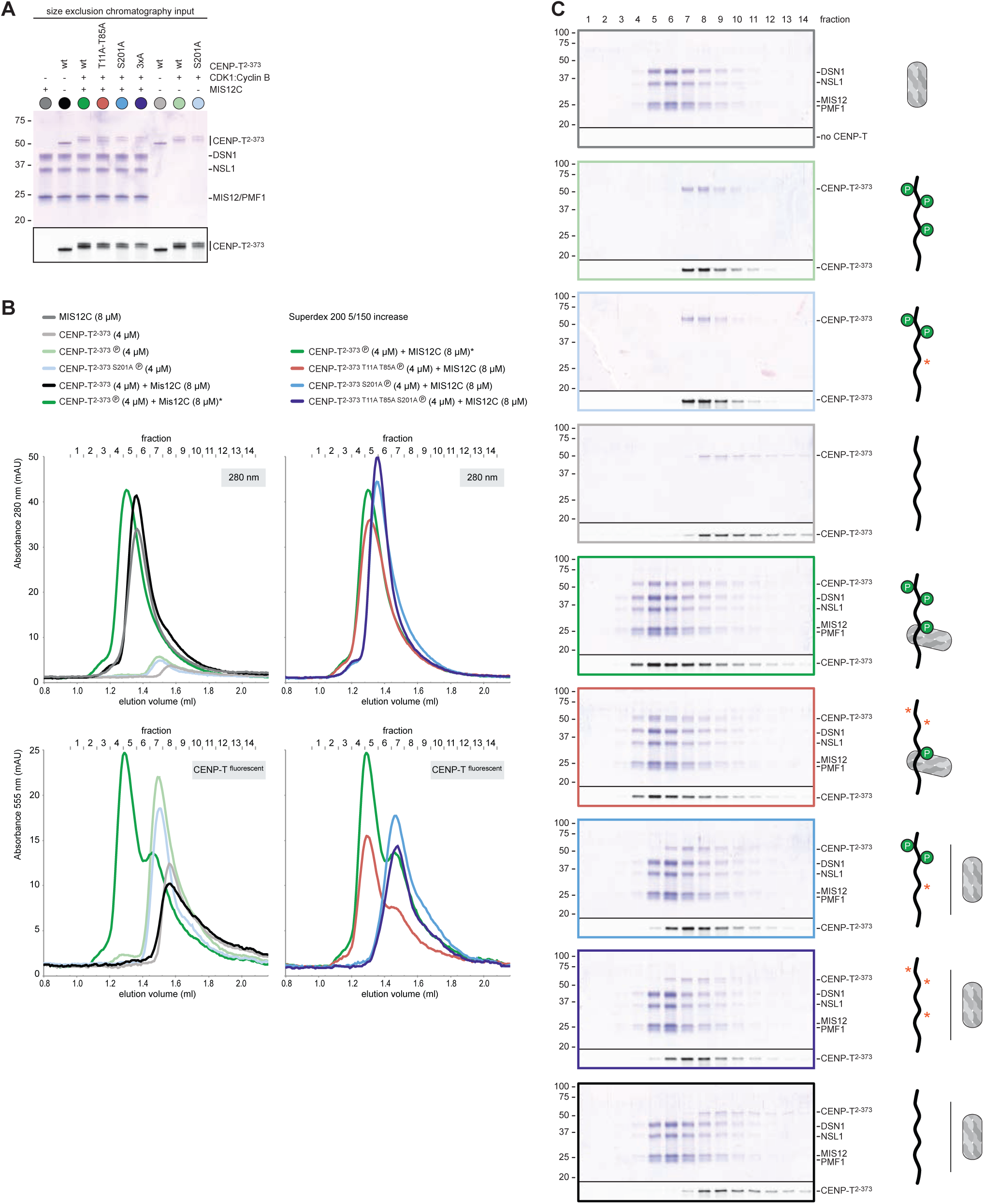
CENP-T phosphorylated by CDK1:Cyclin B at position S201 binds the MIS12 complex. (**A**) A fraction of the mixed samples was analyzed by SDS-PAGE prior to size exclusion chromatography. The phosphorylation-induced shift of CENP-T is clearly visible. (**B, C**) Analytical size-exclusion chromatography (Superdex 200 5/150 increase) and SDS-PAGE show that CENP-T phosphorylated by CDK1 at position S201 binds MIS12C. Data for MIS12C alone (grey), MIS12C mixed with unphophorylated CENP-T (black), and MIS12C mixed with phosphorylated wild type (green), T11A-T85A (red), or S201A (blue) CENP-T are displayed in **Figure 4**.

**Figure 4 - figure supplement 2.**
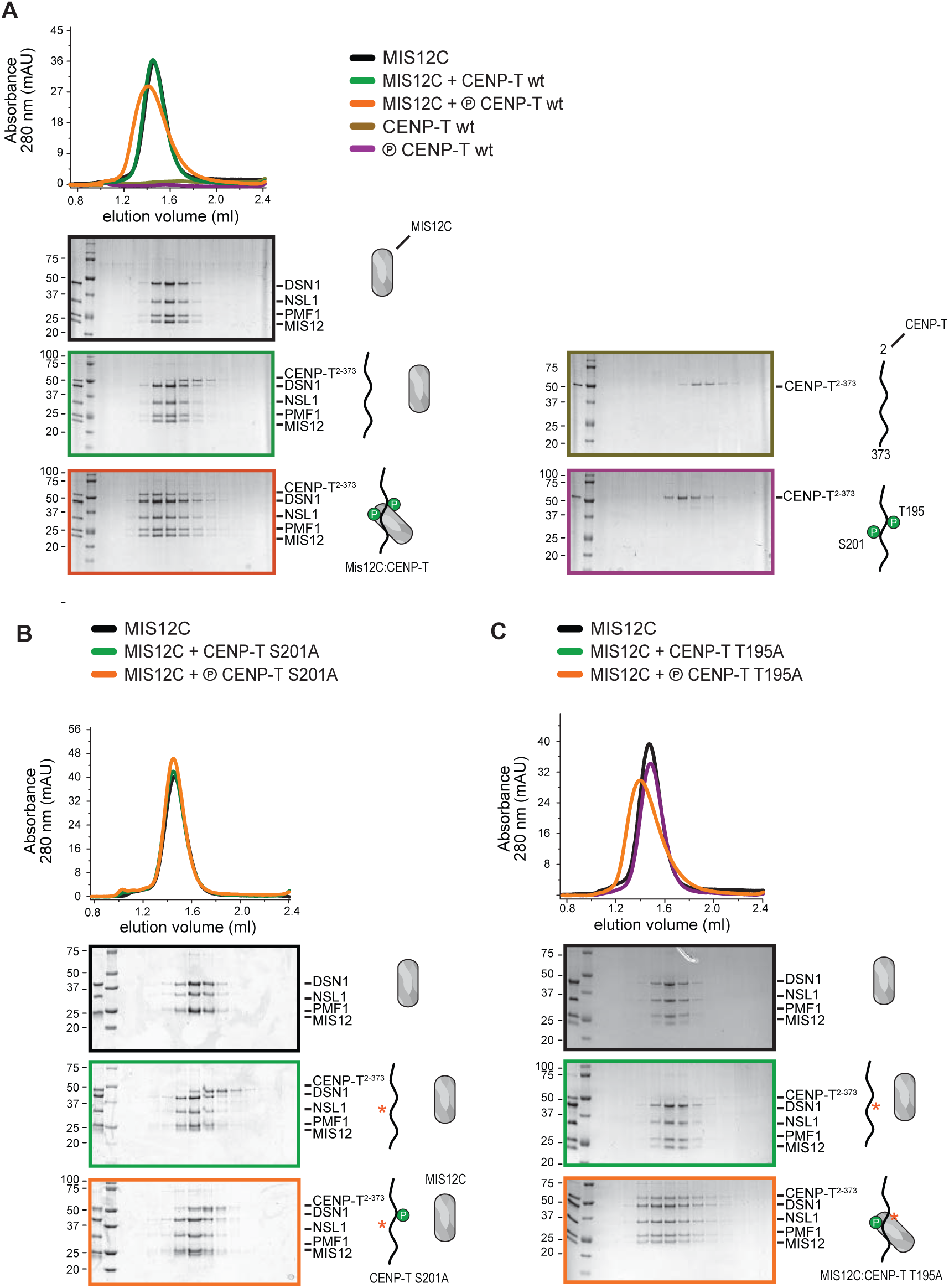
CENP-T mutated at position T195 binds MIS12C. The phosphorylation of CENP-T at position T195 has previously been shown to contribute to the recruitment of MIS12C to CENP-T *in vivo*(Rago et al., 2015). Here we use size-exclusion chromatography (Superdex 200 5/150) and SDS-PAGE to show that phosphorylated CENP-T^T195A^ binds MIS12C *in vitro* whereas CENP-T^S201A^ does not.

**Figure 5 - figure supplement 1.**
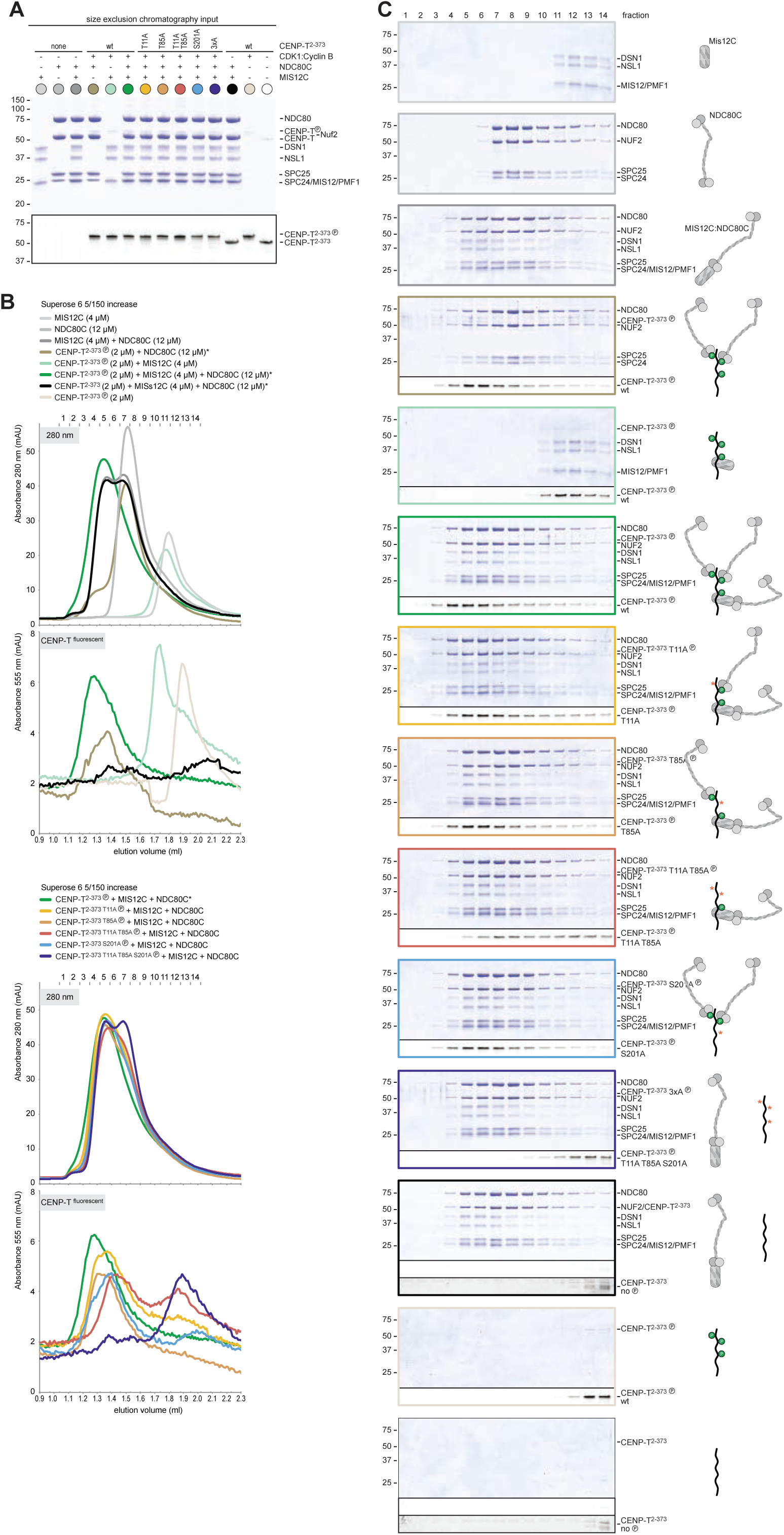
Supercomplex formation depends on the phosphorylation of CENP-T T11, T85, and S201. (**A**) A fraction of the mixed samples was analyzed by SDS-PAGE prior to size exclusion chromatography. The phosphorylation-induced shift of CENP-T is clearly visible, also for the triple CENP-T mutant. (**B**, **C**) Analytical size-exclusion chromatography (Superose 6 5/150 increase) and SDS-PAGE show that phosphorylation by CDK1:Cyclin B of CENP-T at positions T11, T85, and S201 is required for the formation of CENP-T:MIS12C:NDC80C_3_ assemblies. Data for MIS12C:NDC80C alone (grey) and various phosphorylated CENP-T mutants (green, red, blue, purple) are part of **Figure 5**.

**Figure 5 - figure supplement 2.**
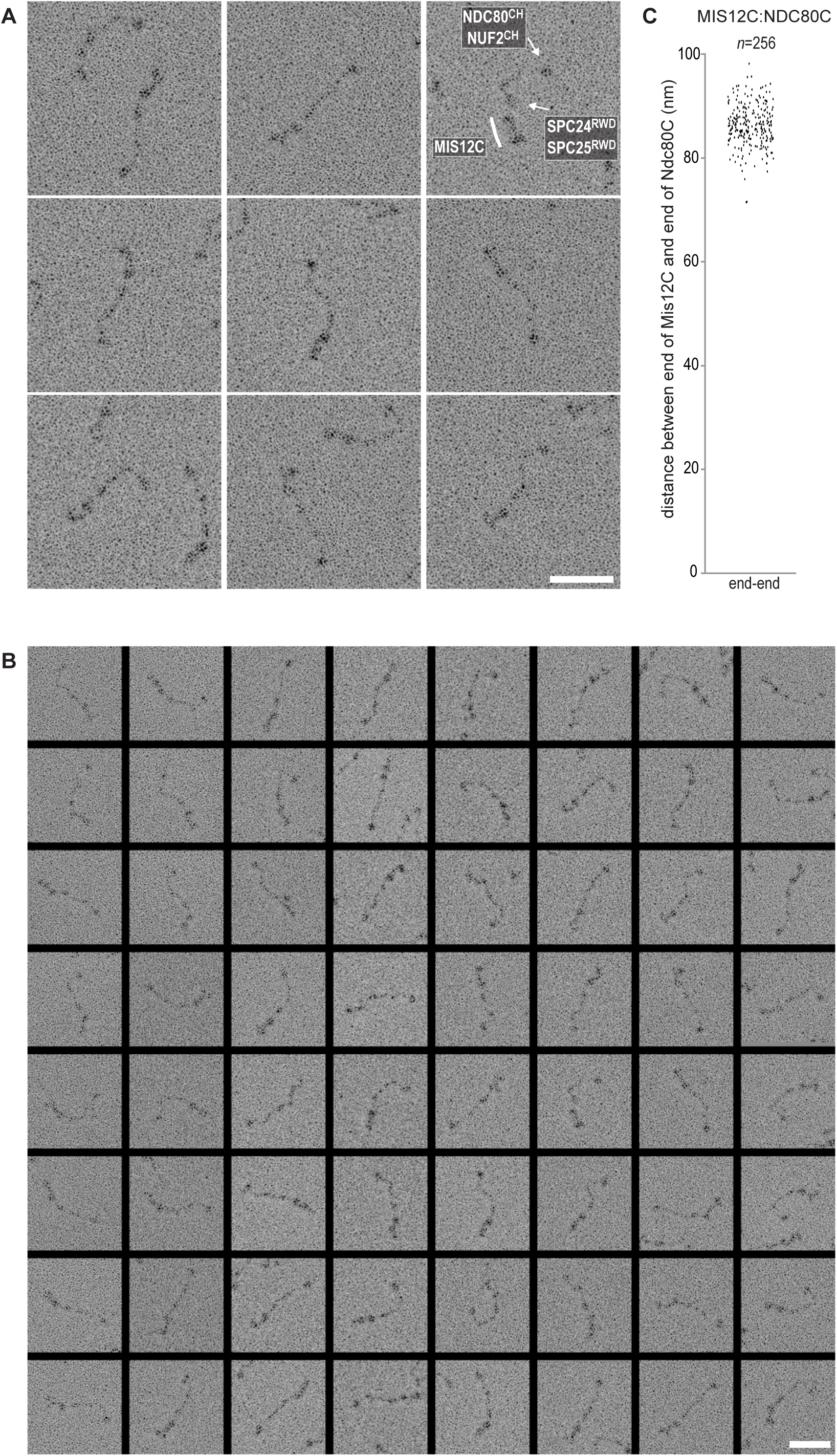
Gallery and measurements of MIS12C:NDC80C. (**A**) MIS12C:NDC80C complexes were visualized after glycerol spraying followed by low-angle metal shadowing. The three micrographs in the top row are also shown in **Figure 5B**. (**B**) Gallery showing 64 MIS12C:NDC80C complexes. Scale bars represent 50 nm. (**C**) End-end and coiled coil distances were measured for the indicated number of shadowed MIS12C:NDC80C complexes. The values shown here are not corrected for an estimated low-angle shadowing contribution of 3 nm at either end.

**Figure 5 - figure supplement 3.**
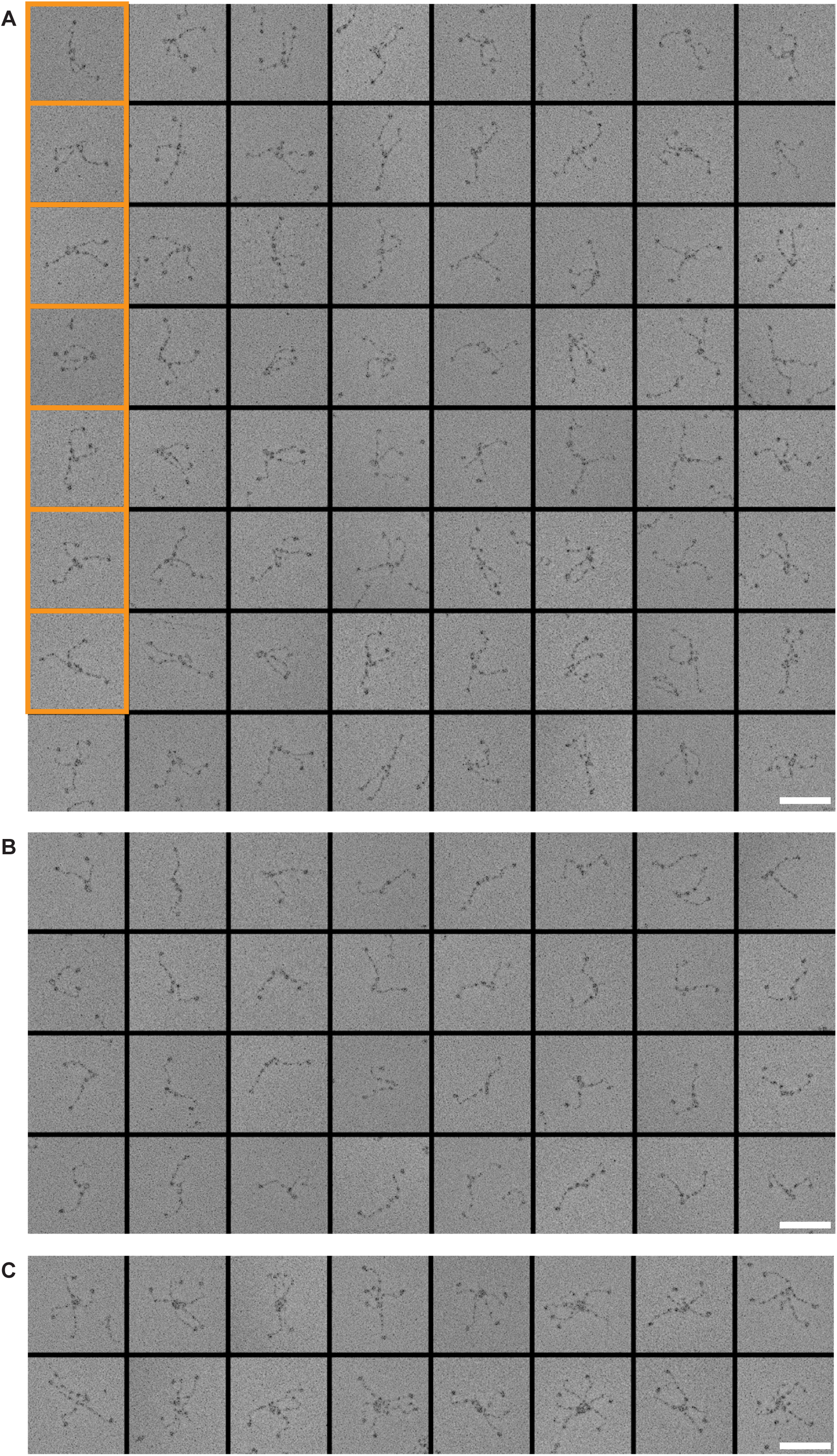
Figure 13 Galleries of CENP-T:MIS12C:NDC80C. CENP-T:MIS12C:NDC80C complexes were visualized after glycerol spraying followed by low-angle metal shadowing. The orange-boxed CENP-T:MIS12C:NDC80C_3_ micrographs are shown in **Figure 5B**. The number of shown CENPT:MIS12C:NDC80C_3_ (panel A; 64), CENP-T:MIS12C:NDC80C _2_(panel B; 32), and CENP-T_x_:MIS12C_x_:NDC80C_>3_ (panel C; 16) micrographs is roughly indicative of the relative abundance of the various assemblies. Scale bars 50 nm.

**Figure 6 - figure supplement 1.**
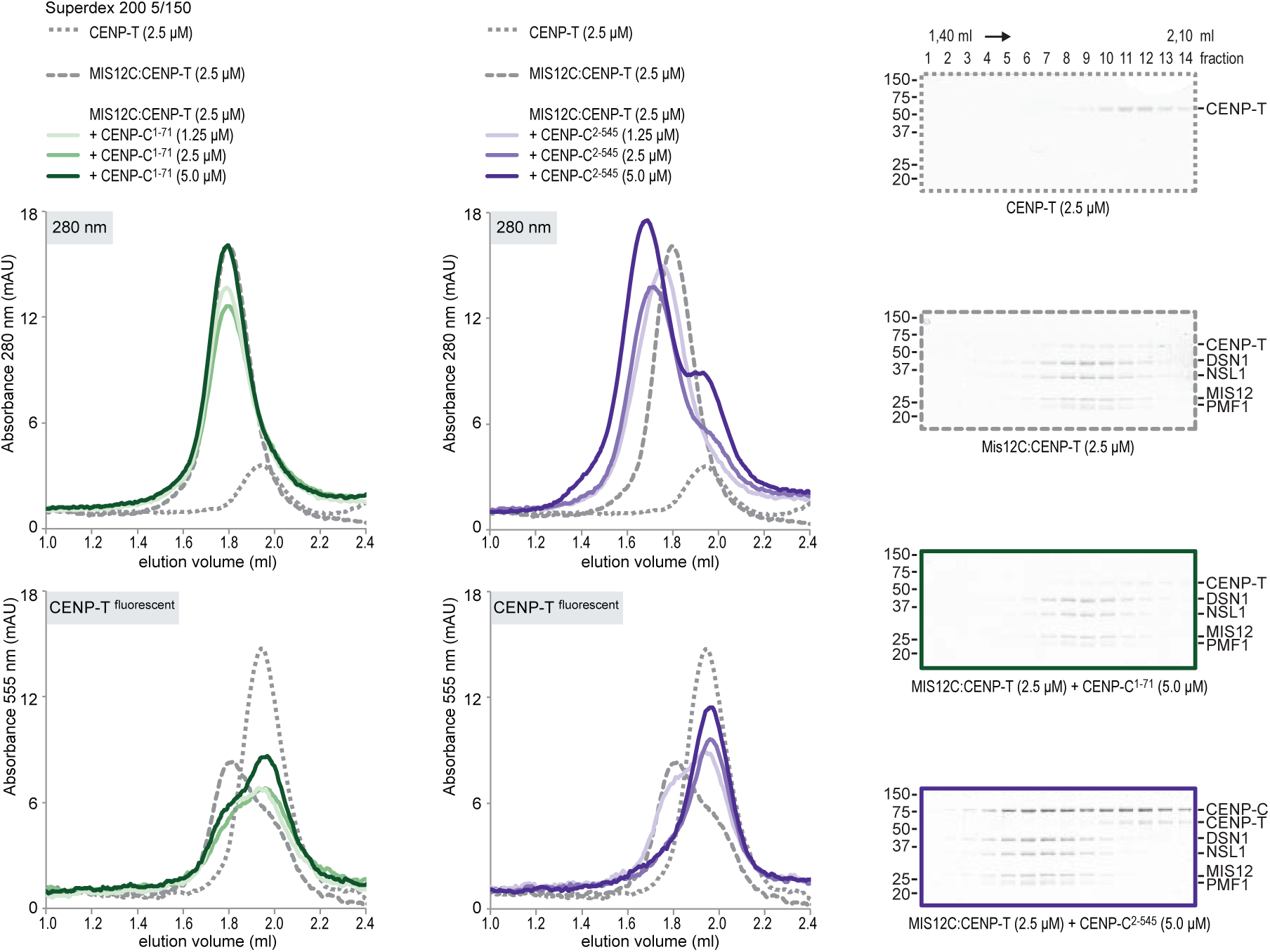
CENP-T and CENP-C are competitive binders of the MIS12 complex. Analytical size-exclusion chromatography (Superdex 200 5/150) and SDS-PAGE show that CENP-C^1-71^ as well as CENP-C^2-545^ form a complex with MIS12C that can no longer bind CENP-T. Fluorescently labeled CENP-T was monitored specifically during chromatography.

